# A sex-specific regulator expands the embryonic ocular field to generate a novel visual system

**DOI:** 10.64898/2026.08.05.742815

**Authors:** Maria Rossello, Tòt Senar-Serra, Rafath Chowdhury, Sophie Tandonnet, Helena García-Castro, Laia Ortega-Flores, Joan Pallarès-Albanell, Marta Gual, Noelle Anderson, Scott W. Roy, Ignacio Maeso, Fernando Casares, Jordi Solana, Isabel Almudi

## Abstract

Morphological novelties often emerge through the redeployment of conserved developmental programs, but how such programs become activated in new developmental contexts remains poorly understood. We investigated the evolutionary origin of the turbanate eyes (TurbEyes) of mayflies, a male-specific additional visual system. Using single-cell transcriptomics, chromatin accessibility profiling and functional genetics, we show that this novelty develops through deployment of the ancestral retinal determination gene network (RDGN). We identify a novel, male-specific paralog of the sex determination transcription factor Doublesex, DsxM, that regulates the eye-specification gene *eyeless/Pax6* and promotes the expansion of the embryonic ocular field. Functional assays impairing *dsxM* activity reduce this male-specific expansion and causes defects in TurbEye development. Our findings reveal a developmental mechanism by which a sex-specific regulator generates a new developmental territory, enabling the spatial redeployment of a conserved developmental program and to generate a novel male-specific visual system.

## Main Text

Key evolutionary transitions have often been associated to morphological novelties. The emergence of new organs in specific lineages often promote their diversification and can result in adaptive advantages which provide opportunities to conquest new ecological niches, such as the wings of insects, the turtle shell, the feathers of birds or the placenta of mammals (*1–4*). Although a few cases in which new organs originated due to the emergence of taxon-restricted genes have been also reported (*5*, *6*), the origin of novel structures often relies on the co-option of conserved pre-existing developmental programs or Gene Regulatory Networks (GRNs) (*3*, *7–9*). Classic examples include butterfly eyespots and beetle horns, which arise through the redeployment of conserved developmental networks in novel developmental contexts (*8*, *9*). In some sexually dimorphic innovations, components of the sex-determination pathway have been implicated in shaping the resulting trait. However, the genetic and genomic changes that trigger GRN co-option, and how sex-specific regulatory mechanisms can direct the deployment of developmental programs to new locations, remain poorly understood.

Despite the profound impact of morphological novelties on insect diversity, the overall organization of the insect head has remained significantly conserved for more than 400 million years (MY)(*10–12*). The insect head comprises a fixed number of segments forming the procephalon, the sensory part, and the gnathocephalon, the feeding structures. The procephalon gives rise to two antennae, two compound eyes, three ocelli and the brain, while the gnathocephalon gives rise to a set of mouth appendages (*13*). Generally, the diversification of insects favored the modification or transformation of pre-existing structures to adapt to different insect lifestyles, rather than addition of new structures. Nevertheless, examples of morphological novelties have been described, particularly in the dorsal region of the head. In most cases, however, these novelties correspond to outgrowths or relatively simple structures, in terms of their morphology or their cell–type composition (*14–16*). In this context, the turbanate eyes (TurbEyes) of mayflies (Ephemeroptera) represent an extraordinary example of morphological novelty (Figure 1A). This novelty is also a striking sexual dimorphism: while females possess the ancestral insect visual system, composed by a pair of lateral compound eyes and three ocelli (Fig. 1B, C), males develop an additional pair of large dorsal compound eyes known as TurbEyes, together with a dedicated set of optic lobes, forming a complete visual system that expands the ancestral insect visual architecture (*17*, *18*)(Fig 1B-C’). Present across Baetidae, the most species-rich family of mayflies with more than one thousand described species, TurbEyes are thought to facilitate the detection of females during the mating swarms (*19*).

**Figure 1.**
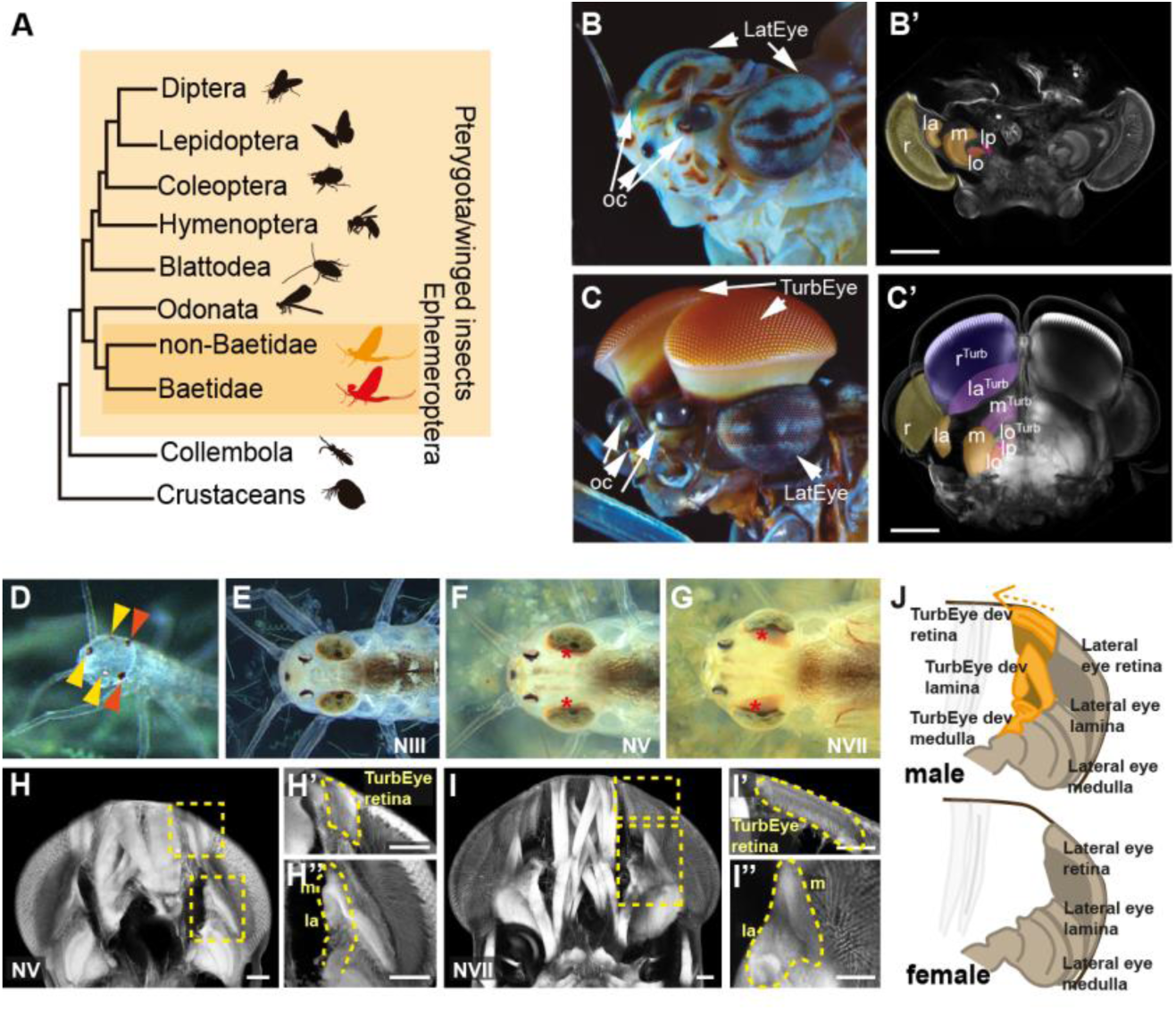
Sexual dimorphism and developmental origin of the mayfly TurbEye. **(A)** Simplified insect phylogeny. **(B, C)** Head morphology of adult females (B) and males (C). Arrows indicate the lateral eye (LatEye) and the male-specific TurbEye. White arrowheads denote the dorsal expansion characteristic of the TurbEye. oc, ocelli. **(B’, C’)** Cleared heads of female (B’) and male (C’); r, retina; la, lamina; m, medulla; rTub, laTub and mTub correspond to TurbEye-specific visual neuropils. **(D–G)** Dorsal views of nymphal development showing the ocelli (yellow arrowheads) and lateral compound eyes (red arrowheads) in hatchlings (D) and the emergence of the TurbEye primordia (asterisks, F, G). Developmental stages are indicated in each panel. **(H, I)** Cleared male heads at nymph V (H) and nymph VII (I). Insets show higher magnification of the developing TurbEye region. **(J)** Schematic representation of visual-system organization in females and males. Scale bars: 50 μm.

The ontogeny of this set of male TurbEyes is temporally de-coupled from those of the lateral compound eyes and ocelli, which are fully formed and functional when nymphs hatch (Fig. 1D). By contrast, a major part of TurbEye development occurs during nymphal stages, in which a pair of bilayered epithelial primordia grow towards the medial part of the head, together with their associated optic lobe regions (lamina, medulla and lobula; Fig 1E-J). Despite the remarkable nature of the TurbEyes, the GRNs and developmental mechanisms responsible for the development and origin of this morphological novelty remain unexplored.

Here, we study the genetic mechanisms underlying the evolution of the TurbEye in the Baetidae mayfly *Cloeon dipterum* and describe how a novel paralog of the sex-determination gene *doublesex* (*dsx*), enables the male-specific deployment of the ancestral retinal developmental program through the regulation of *eyeless* (*ey*), generating a novel visual system.

### The turbanate eye of males develops during nymphal stages through the co-option of the RDGN

To elucidate the genetic mechanisms underlying the development of the TurbEyes, we performed scRNA-seq on heads of three nymphal stages: NIII, in which the TurbEye primordia are still not externally detectable, NV, in which TurbEye primordia start to be visible and NVII, when the primordia have acquired half of the final size of the TurbEye retinas (Figure 1E-G). We generated specific male and female samples for stages NV and NVII, due to the presence of the TurbEye primordia, whereas samples from NIII were a mixture of female and male heads, since the absence of such primordia precluded us from distinguishing the sexes at the time of sample collection. To minimize dissociation-induced artefacts and improve reproducibility in single-cell transcriptomics (*20*), we leveraged ACME fixation (*21*) and SPLiT-seq combinatorial barcoding (*22*) to profile two biological replicates from each of the five experimental conditions (see Methods, Table S1 and Fig. S1). Cell clustering using Leiden algorithm (*23*) yielded 32 cell clusters (Fig 2A-B, Fig S2). To annotate these clusters, we used the five highest-ranked marker genes identified by differential expression analysis (Wilcoxon rank-sum test (*24*); Table S2) together with known marker genes inferred from orthology to *D. melanogaster* genes (Table S3). These main cell clusters included cell types or states belonging to the main cell types expected in nymphal heads of mayflies, such as muscle cell populations (Cluster (Cl) 3), glia cells (Cl 12 and 24), seven epidermal or cuticular cell clusters (Cl 4, 6, 7, 8, 17, 18, and 20) and seven annotated cell types related to visual systems (Cl 0, 1, 2, 9, 11, 13, 19; Fig. 2A-B, Fig. S2).

**Figure 2.**
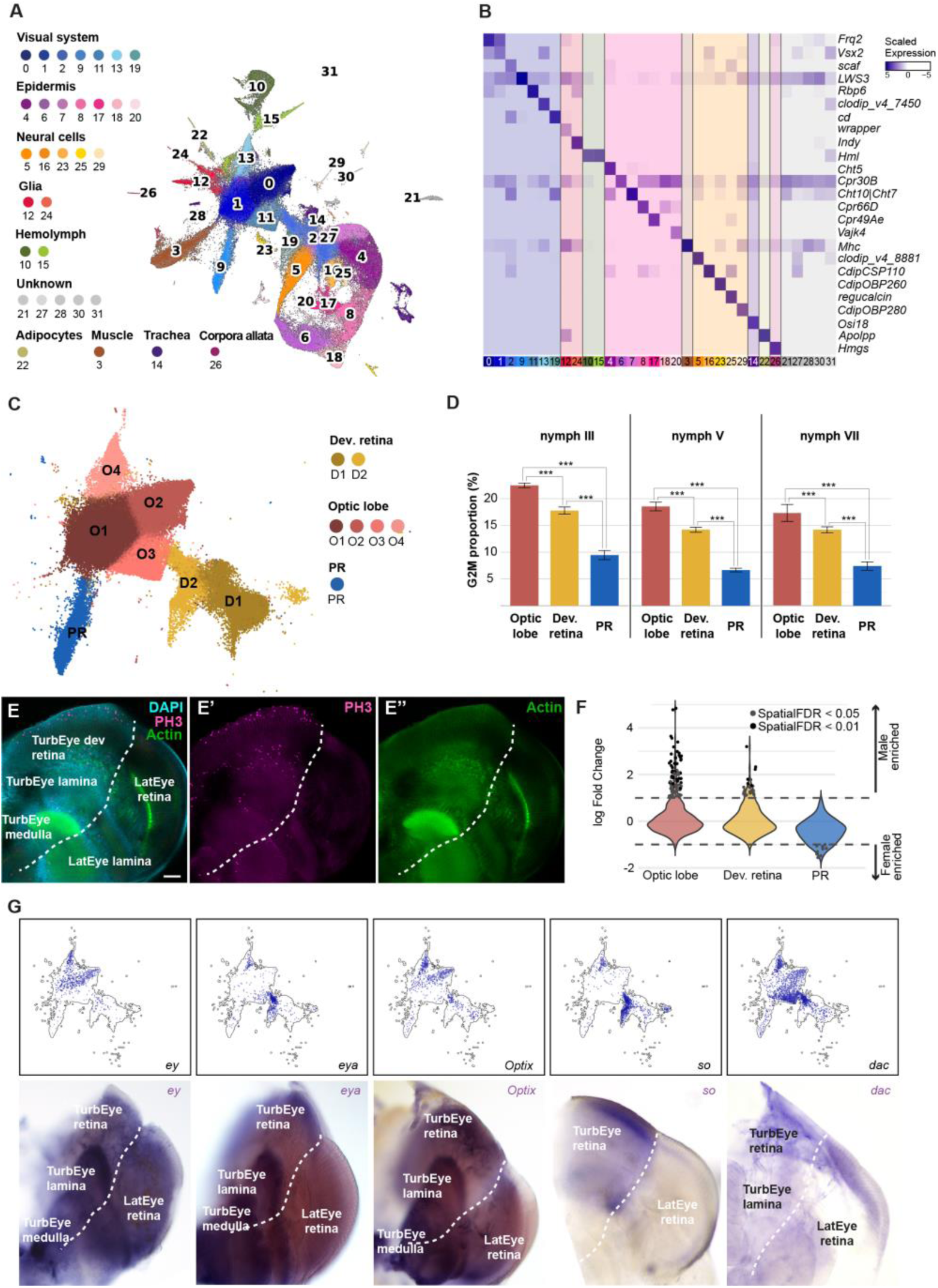
Single-cell characterization of the developing visual system. **(A)** UMAP projection of single-cell transcriptomes from developing male heads showing major cell populations. **(B)** Heatmap of representative marker genes across cell clusters. **(C)** Subclustering of visual-system cells identifying optic lobe (O1-O4), developing retina (D1-D2), and photoreceptor (PR) populations. **(D)** Proportion of G2/M-phase cells in visual-system populations across developmental stages. **(E–E″)** Confocal images of the developing visual system stained with DAPI, PH3, and Actin. Dashed lines indicate tissue boundaries. Scale bar: 50 μm **(F)** Distribution of sex-biased expression in visual-system cell populations. **(G)** Expression of selected retinal determination genes visualized in the single-cell dataset (top) and by in situ hybridization (bottom).

Next, we performed subclustering on these visual-system-related cell clusters to resolve the cellular composition of the developing visual system. This sub-clustering revealed seven subpopulations that grouped into three major categories, supported by PAGA analysis (*25*): a photoreceptor cluster (PR), two developing retina clusters (D1 and D2) and four optic lobe clusters (O1-O4; Fig. 2C, Fig. S3, Table S4). Cell identities were assigned based on the expression of diagnostic markers and their correspondence with known components of the insect visual system (Fig. S4; Table S4). The PR cluster was characterized by high expression of opsin genes and likely corresponds to differentiated photoreceptors from the lateral compound eyes and ocelli. In contrast, TurbEye photoreceptors were still immature at the nymphal stages analyzed in our scRNA-seq experiment and did not yet express opsins (Fig. S5), suggesting that cells of the nascent TurbEye were instead represented within the developing-retina populations (D1 and D2). O1–O4 corresponded to distinct optic-lobe populations (Fig. 2C, Fig. S4). Analysis of cell-cycle states revealed that both the developing-retina and optic-lobe populations contained substantially higher proportions of G2/M-phase cells than differentiated photoreceptors across all three nymphal stages, indicating sustained proliferative activity in the visual-system progenitors (Fig. 2D, Fig. S6). Consistent with these observations, Phospho-histone H3 (PH3) immunostaining identified larger number of mitotic cells in the regions that will give rise to the TurbEye visual system (Fig 2E). To characterize whether these visual system populations exhibited sex-bias (i.e., some clusters were enriched in cells belonging to male or female samples), we performed differential abundance analysis with Milo (*26*). We observed a higher representation of male cells in the optic lobe and developing retina (cl O1, O2, O3, O4 and D2; Fig. 2F, Fig. S7), which was consistent with the increase of male cells due to the growth and differentiation of the TurbEye primordia and their associated optic lobes.

The Retinal Determination Gene Network (RDGN) is a highly conserved GRN which is responsible of the development of eyes in very distantly related animal lineages (*27–29*). Consistent with the deployment of a canonical retinal developmental program, all major RDGN transcription factors (TFs), including *eyeless/Pax6 (ey), sine oculis/Six1,2 (so), optix/Six3,6, dachshund (dac)*, and the tyrosine-threonine phosphatase and transcriptional co-activator *eyes absent (eya)*, were expressed in the developing TurbEye and its associated optic lobes (Fig. 2G). These expression patterns indicate that the TurbEye does not arise through a lineage-specific GRN, but rather through the redeployment of the ancestral RDGN. However, how this conserved developmental program became specifically activated in the male TurbEye primordium remained unknown.

### A novel male-specific Doublesex paralog emerges as potential regulator of TurbEye development

To identify upstream regulators responsible for activating the retinal developmental program in the male TurbEye, we performed differential expression analyses between male and female samples using pseudobulk profiles from the developing retina and optic-lobe populations. This analysis revealed numerous sex-biased genes enriched in males, including several TFs (Fig. 3A, Fig. S8, Table S5-S6, see methods). Among them was a 125 amino acid (aa) predicted TF with a DMRT-type DNA binding domain that we identified through phylogenetic analysis as a novel third paralog of the insect sex determinant Doublesex (Dsx) (*30–32*), (Fig 3A, Fig. S9). PCR amplification from genomic DNA detected this gene exclusively in males, whereas no amplification was observed in females (Fig. S9). We therefore named this paralog *dsxM.* Strikingly, *dsxM* was among the most strongly male-enriched TFs in both the developing-retina and optic-lobe populations (Fig. 3A, Fig. S8), suggesting a putative role in the development of both, the neuroepithelium that will give rise to the TurbEye retina and its associated optic lobe structures. Consistent with a specialized function in TurbEye development, *dsxM* showed little evidence of expression outside the visual-system and epidermal populations (Fig. S8). In situ hybridization of this TF in nymphal heads revealed that *dsxM* was expressed in both, the retinal and the optic lobe regions of the developing TurbEye, while absent from the lateral eye and its optic lobe (Fig 3B).

**Figure 3.**
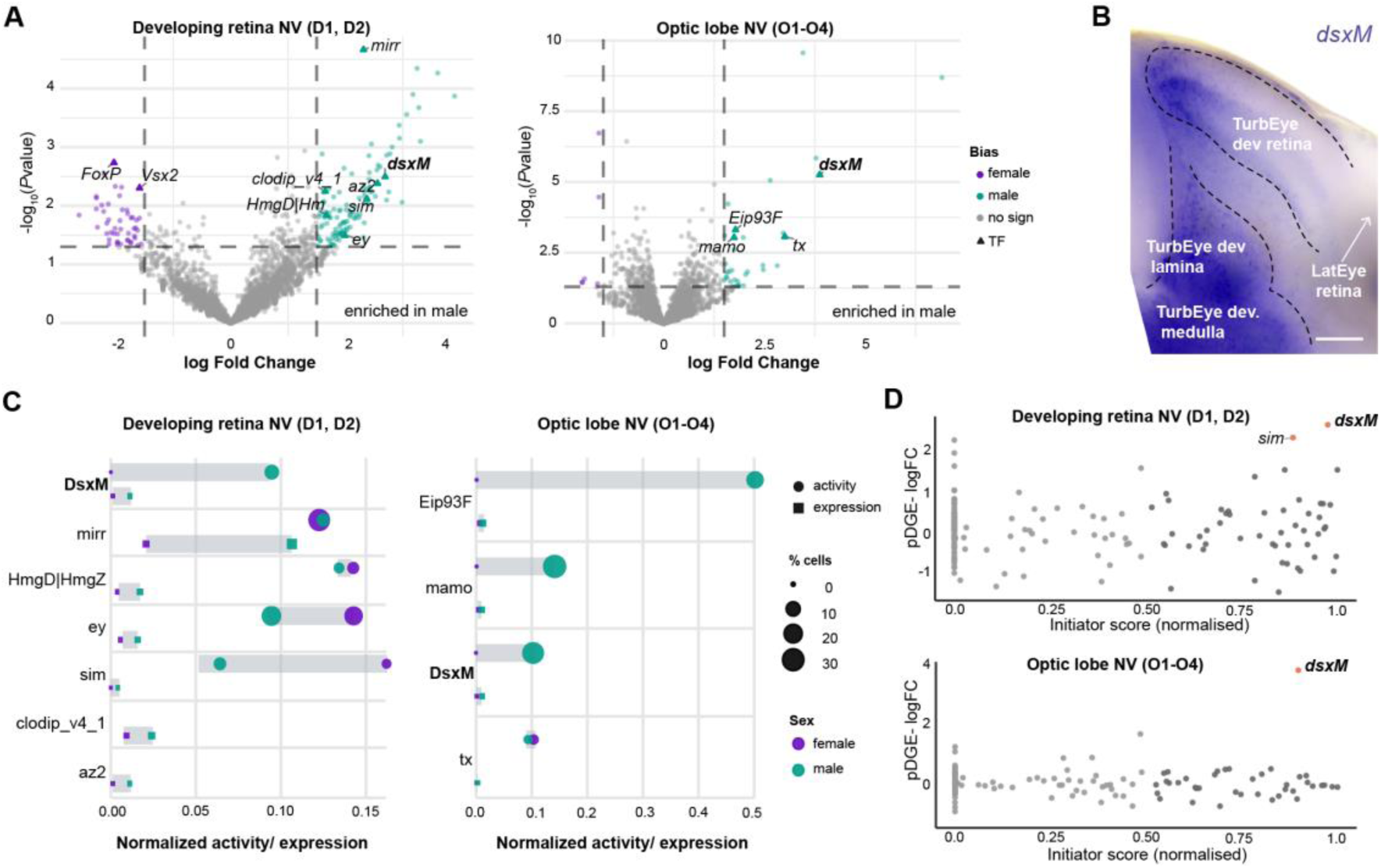
Male-biased expression and regulatory activity of *dsxM* in developing visual tissues. **(A)** Volcano plots showing sex-biased gene expression in the developing retina (left) and optic lobe (right) at nymph V, with dashed lines marking the significance thresholds (P < 0.05, |logFC| ≥ 1.5). Genes passing these thresholds are coloured in teal (male-biased) or purple (female-biased), with significant transcription factors highlighted as larger, labelled triangles. **(B)** In situ hybridization showing *dsxM* expression in the developing male visual system, with dashed lines outlining the TurbEye-specific developing retina, lamina, and medulla, and the lateral eye (LatEye) retina. Scale bar: 50 μm. **(C)** Normalised regulon activity (normalized SCENIC AUC) and pseudobulk normalized expression of male differentially expressed transcription factors in developing retina (left) and optic lobe (right), by sex; dot size indicates the percentage of cells in which each factor is active or expressed. **(D)** Scatter plot showing normalised initiator score against pDGE logFC in developing retina (top) and optic lobe (bottom). Genes in light grey pass neither the initiator score nor the fold-change threshold; genes in dark grey pass the initiator score threshold but not the fold-change threshold; genes in orange pass both thresholds.

To further evaluate the regulatory activity of the candidate transcription factors identified in the differential expression analyses, we integrated our scRNA-seq data with ATAC-seq datasets generated from matching developmental stages using a customized SCENIC pipeline (*33*) (see methods and extended methods, Fig. S10, S11, Dataset S1, Dataset S2, Table S7, Table S8). This approach reconstructs gene regulatory networks and identifies regulons, defined as transcription factors and their predicted target genes, allowing the activity of each regulon to be quantified at single-cell resolution. Comparison of regulon activity between male and female cells revealed several male-biased regulons in both the neuroepithelium and the optic lobe, including regulons associated with *mirror* (*mirr*)*, HmgD/HmgZ, mamo* and *Eip93F* (Fig 3C, Figure S12, Table S9, see methods and extended methods). Notably, the *dsxM* regulon was the only one that displayed elevated activity in male cells in both structures of the visual system. Finally, to prioritize candidate upstream regulators, we calculated an initiator score, a network-based metric that identifies transcription factors located near the top of the inferred regulatory hierarchy (see methods and extended methods, Fig. S13, Table S10, Table S11). Plotting initiator score against sex-biased expression revealed *dsxM* as the most prominent candidate in both neuroepithelial and optic-lobe populations (Fig. 3D).

### DsxM promotes the expansion of the embryonic *ey* domain and TurbEye development

To identify the earliest stage at which dsxM may function during TurbEye development, we decided to investigate the onset of *dsxM* expression. Analysis of previously generated embryonic RNA-seq datasets (*34*, *35*) revealed that *dsxM* expression begins during stages 6-8 of embryogenesis, coincident with head regionalization and overlaps temporally with several head-patterning and retinal determination genes, including *ey, wg, oc* and *Optix* (Fig. 4A). These observations suggested that *dsxM* might act early during head development, before the onset of TurbEye morphogenesis.

**Figure 4.**
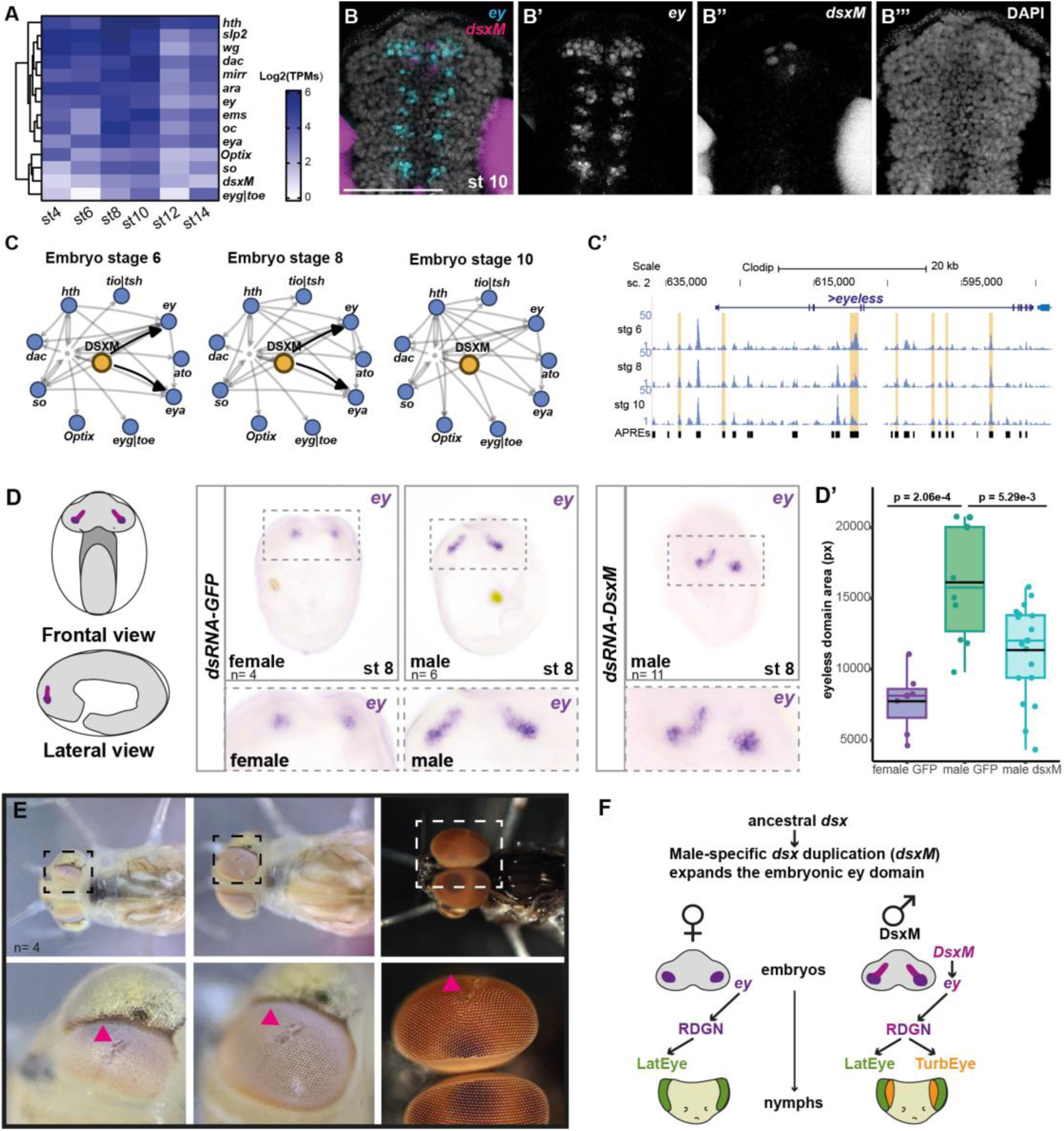
DsxM promotes expansion of the embryonic ocular field and TurbEye formation**. (A)** Expression dynamics of retinal determination genes and *dsxM* during embryogenesis. **(B-B’’’)** Hybridization Chain Reaction (HCR) showing expression of *ey* (B’) and *dsxM* (B’’) in the ocular primordia of a male embryo at stage 10. DAPI (B’’’). **(C)** Inferred gene regulatory networks across embryonic stages 6, 8 and 10. Black arrows indicate predicted direct regulatory interactions from DsxM to RDGN components. Grey edges represent additional inferred interactions within the network. **(C’)** Chromatin accessibility landscape of the *eyeless* locus during embryonic development. Accessible regions containing predicted Dsx-binding motifs are highlighted in orange. APREs, Accessible Putative Regulatory Elements. **(D)** *ey* expression in control (dsRNA-GFP) and *dsxM* RNAi stage 8 embryos. Insets show higher magnification of the ocular primordia. Sample sizes are indicated in each panel. **(D’)** Quantification of the *ey* expression domain area in female control, male control and male *dsxM* RNAi embryos. p-values from Wilcoxon test are shown above comparisons. **(E)** Representative F0 mosaic *dsxM* CRISPR individual exhibiting localized defects in TurbEye morphology during nymphal and adult stages (arrowheads). **(F)** Proposed model for the role of the male-specific *doublesex* duplicate (*dsxM*) in TurbEye origin. Scale bar: 50 μm.

We next focused on *ey*, a RDGN gene capable of triggering the deployment of the eye developmental program, and assessed its spatial relationship with *dsxM*. HCR experiments revealed partial overlap between both genes in the dorsal procephalon, a region that contributes to the ocular territory of the head (*12*)(Fig. 4B). This overlap suggested a potential regulatory interaction occurring during embryonic head patterning. To further explore this possibility, we integrated previously generated embryonic RNA-seq and ATAC-seq datasets (*35*) and reconstructed stage-specific regulatory networks, using ANANSE (*36*)(Fig. S14, Dataset S3, see methods and extended methods). The GRNs reconstructed in these embryonic stages predicted that DsxM is directly activating the expression of *ey* during st 6 and st 8 of *C. dipterum* embryogenesis (Fig 4C, Fig. S15). In agreement with this prediction, several accessible regions within the *ey* locus contained predicted Dsx-binding motifs, suggesting that DsxM may regulate *ey* through direct interaction with its *cis*-regulatory landscape (Fig. 4C’, Table S12). To confirm this, we established gene knock-downs through parental RNAi (i.e. injection of dsRNA into the abdomen of gravid females, see methods), since *C. dipterum* is one of the few ovoviviparous species of mayflies (*37*, *38*). Gravid females were injected with dsRNA against *dsxM* at st 6 of embryogenesis to avoid earlier deleterious effects and fixed the embryos at st 8 to analyze the phenotypes. As controls, gravid females were injected with dsRNA against the Green Fluorescent Protein (GFP) and processed identically (see methods). In control embryos, the *ey* expression domain occupied a significantly larger area in males than in females (Fig. 4D-D’, Fig. S16). Knockdown of *dsxM* resulted in a significant reduction of this male-specific expansion, resulting in *ey* domains substantially smaller than those of control males and, in some individuals, approaching the dimensions observed in females (Fig. 4D-D’, Fig. S16). Together, these results indicate that DsxM regulates *ey* expression and is required for the establishment of the expanded male *ey* domain during embryogenesis, supporting a role for DsxM in generating the enlarged ocular field that precedes TurbEye development.

Because embryonic *dsxM* knockdown prevented us from assessing the developmental consequences of the altered *ey* domain at later stages of TurbEye formation, we next generated mosaic loss-of-function individuals using DIPA-CRISPR (*39*)(see Methods). Mosaic individuals carrying localized disruptions of *dsxM* displayed clear defects in the TurbEye, including regions with abnormal ommatidial organization adjacent to apparently unaffected tissue (Fig. 4E, Fig. S17). The spatial restriction of these defects is consistent with the mosaic nature of the mutagenesis. Together with the persistent expression of dsxM during nymphal development, these phenotypes suggest that dsxM contributes not only to the establishment of the embryonic ocular field but also to subsequent stages of TurbEye morphogenesis. Taken together, these results identified DsxM as a central regulator of TurbEye development in *C. dipterum* and supported a model in which the male-specific expansion of the embryonic *ey* domain contributes to the formation of this evolutionary novelty (Fig. 4F).

## Discussion

The origin of morphological novelties remains a central question in evolutionary developmental biology, particularly in cases where novel structures arise within otherwise conserved body plans, such as the insect head (*15*). Our results show that the TurbEye of the mayfly *C. dipterum* develops through the deployment of the canonical retinal determination gene network (RDGN) into a novel developmental territory. We identify a male-specific paralog of *doublesex* (*dsxM*) as a central regulator of this process, acting upstream of *ey* to promote the expansion of the embryonic ocular field and the formation of the male-specific visual system (Fig. 4F). Hence, our findings reveal a developmental mechanism through which a sex-specific regulator can expand a pre-existing developmental field, enabling the spatial redeployment of an ancestral developmental program in a novel context.

Co-option of GRNs is a common mechanism underlying morphological novelty (*40*, *41*). However, the genetic and developmental changes that initiate these co-option events remain far less understood. Our results support that the TurbEye develops through the deployment of the canonical RDGN, including all major components of this deeply conserved network (Fig 2). Similar patterns of incomplete co-option and subsequent specialization have been documented in other morphological innovations, where ancestral GRNs are co-opted and subsequently modified to generate lineage-specific morphologies (*42*, *43*). In the case of the TurbEye, our findings suggest that activation of a single upstream regulator, *ey*, in a novel cephalic location may be sufficient to trigger deployment of the entire retinal developmental program, as *ey/Pax6* can induce eye development in non-ocular tissues in both *Drosophila* and *Xenopus* (*27*, *44*). These findings point to expansion of a pre-existing developmental field as a developmental route through which conserved GRNs can be deployed in novel spatial contexts.

The identification of a new male-specific paralog of Dsx as a key TF involved in the TurbEye development aligns with previous work identifying Dsx as a central regulator of sexually dimorphic novelties, including the horns of horned beetles (*45*, *46*). These studies have shown that Dsx can function as a regulatory hub, able to acquire novel target genes and to mediate tissue-specific developmental programs through changes in cis-regulatory regions. In contrast to previously described Dsx targets involved in the differentiation of sexually dimorphic traits (*45*, *47*), our results indicate that DsxM regulates *ey*, a master regulator of ocular specification, revealing a previously unknown connection between the sex-determination pathway and the retinal determination network. This finding suggests that sex-determination pathways can influence morphological innovation through interactions with deeply conserved organ-specification networks.

The TurbEye is particularly remarkable in the context of insect head evolution. While evolutionary novelties of the insect head are relatively uncommon and often involve structures of limited cellular complexity (*15*, *16*), the TurbEye represents the emergence of an additional sensory system within an otherwise highly conserved body plan. This visual system is more than a simple morphological novelty. In addition to comprising an extra retina and associated optic lobes, previous work has shown that it is associated with the recruitment of TurbEye-specific opsins and likely contributes to male-specific visual behaviors (*19*, *38*). Thus, the spatial redeployment of the RDGN appears to have generated a novel developmental territory that subsequently diversified structurally and functionally. Rather than producing a mere duplicate of the ancestral visual system, this process gave rise to a specialized visual organ whose morphology, gene expression profile and likely behavioral role differ from those of the lateral compound eyes. Together, our findings provide a mechanistic example of how a conserved developmental program can be redeployed in a novel location during evolution. The TurbEye illustrates how the generation of a new developmental territory can create opportunities for subsequent morphological and functional diversification.

## ACKNOWLEDGMENTS

We thank R. Hedley at the Flow Cytometry Facility at the Dunn School of Pathology (University of Oxford) and the Optical Microscopy Unit of the CCiT of the Universitat de Barcelona. We are also very thankful to Alberto Perez-Posada, David Salamanca, Marta Portela, Alba Almazán and the Solana and Almudi labs for experimental and bioinformatics advice. We also thank Manuel Irimia and Marta Iglesias for critically reading the manuscript.

## Funding

This research was funded by the HORIZON EUROPE European Research Council research and innovation program (ERC-CoG2021-101043751 to I.A.), by the Ministerio de Ciencia, Innovación y Universidades (PID2020-116041GB-I00 and PID2023-151401NB-I00 to I.A.; PGC2018-093704-B-I00 to F.C.) and by the Ministerio de Ciencia, Innovación y Universidades and the European Commission NextGenerationEU/PRTR program (CNS2023-145403). M.R. was awarded a Margarita Salas fellowship from Ministerio de Ciencia e Innovación. ST holds a HORIZON-MSCA-2023-PF-01 (101147787).

## Author contributions

Conceptualization: I.A., J.S., M.R., T.S.-S., F.C.; Methodology: M.R., T.S.-S., S.T., R.C., H.G.-C., L.O.-F., J.P.-A., I.A.; Investigation: M.R., T.S.-S., S.T., R.C., H.G.- C., L.O.-F., M.G., I.M., I.A.; Visualization: M.R., T.S.-S., R.C., J.P.-A., I.A.; Funding acquisition: I.A., J.S., FC; Project administration: I.A.; Supervision: I.A., J.S.; Writing – original draft: I.A., M.R.; Writing – review & editing: all authors.

## Competing interests

The authors declare that they have no competing interests.

## Materials and methods

### Culture maintenance and sample collection

Samples were obtained from a *C. dipterum* culture maintained in the laboratory as previously described in (*38*). Embryos at different developmental stages were collected by dissecting gravid females that had been fertilized on different days. Nymphs at the desired stages were selected, and their heads were dissected for subsequent processing.

### Single-cell RNA-seq library preparation

Single-cell transcriptome profiling was performed using an adapted SPLiT-seq protocol (*22*) as described in (*48*) and optimised for *C. dipterum*. Heads were dissected from developmental stages NIII, NV, and NVII (both sexes, two biological replicates per condition) under a stereomicroscope and dissociated following the ACME fixation–dissociation method (*21*), which preserves RNA integrity during cell isolation. Further description of the SPLiT-seq protocol and library preparation are described in extended methods.

Sequencing was performed on an Illumina NovaSeq 6000 platform (Novogene) using paired-end reads according to the SPLiT-seq design, with sequencing depth chosen to achieve sufficient coverage per cell for downstream analysis.

### Single-cell RNA-seq computational processing and quality control

Raw sequencing data were processed using BarQC (*49*), a tool for demultiplexing, barcode correction, and quality control of SPLiT-seq data (Table S1). Reads were trimmed to remove adaptors and poly(A) tails, aligned to the aligned to the *Cloeon dipterum* reference genome (GCA_902829235.1) using clodip_v4 gene models (*50*), and merged with corresponding cell barcode information. Gene annotations were added, and digital gene expression matrices were generated after applying cell- and gene-level filtering thresholds.

For the resulting matrix, filtering thresholds for minimum counts, minimum genes detected, and maximum mitochondrial fraction were optimised empirically by inspecting knee-point distributions and quality metric scatter plots. Highly variable genes (HVGs) were identified using cell_ranger (*51*), and the number of principal components retained for neighbourhood graph construction was determined by elbow-plot inspection. Clustering was performed with the Leiden (*23*) algorithm across a range of resolutions, and the resolution yielding biologically interpretable, non-redundant clusters was selected.

Cluster marker genes were identified by differential expression analysis in Scanpy (*52*), using the Wilcoxon rank-sum test with Benjamini–Hochberg correction (Table S2). Clusters were annotated to cell types based on the expression of known *Cloeon dipterum* marker genes and orthology to *Drosophila melanogaster* reference cell types (Table S3). To characterize the visual system in greater resolution, eye-related clusters were extracted and subjected to iterative subclustering, applying the same HVG selection, dimensionality reduction, and clustering workflow to the subset (Table S4).

Differential cell-type abundance between male and female samples within the visual system was assessed using Milo (*26*), which frames abundance testing in terms of overlapping k-nearest-neighbour (KNN) neighbourhoods to account for the discrete nature of clusters and the continuous structure of transcriptional space. This analysis identified cell populations significantly enriched in males, providing a cell-type-level view of sexual dimorphism in the developing visual system.

Finally, cell cycle phase was assigned to each cell by scoring the expression of S-phase and G2/M-phase gene sets derived from (*53*). Cell cycle phase proportions were compared between male and female cells within each visual system population using binomial generalized linear mixed models (GLMMs), fitted with biological replicate as a random effect to account for inter-replicate variability (lme4; (*54*)). Overall sex effects were assessed by Type III likelihood ratio tests, and pairwise comparisons between groups were performed using estimated marginal means with Tukey adjustment (*55*).

### Transcription factor identification

*Cloeon dipterum* TF candidates were identified using a multi-evidence pipeline combining protein domain annotation (InterProScan against Pfam/AnimalTFDB), GO-term filtering (structure- and domain-based), and orthology/homology to curated Drosophila melanogaster TF and cofactor lists. Candidates were selected by the number of independent evidence levels supporting them and flagged for cofactor activity (Table S13). For more information, see extended methods.

### Pseudo-bulk differential gene expression

Sex-biased gene expression was assessed by pseudo-bulk differential expression (pDGE) on the scRNA-seq data. Cells were aggregated into pseudo-bulk samples by sex, developmental stage, and replicate, and, where relevant, by cell type, requiring genes to be expressed in at least 5 cells per condition. Counts were filtered, TMM-normalised, and analysed with limma-voom (*56*, *57*), testing male vs. female contrasts within each developmental stage. Genes were called differentially expressed at pvalue < 0.05 and |logFC| ≥ 1.5.

### ATAC-seq library preparation

ATAC-seq from (*58*) was optimized for *C. dipterum* as described in (*35*) and applied here to dissected heads. Briefly, heads from NIV, NVI, and NVII developmental stages (both sexes, two biological replicates per condition) were homogenized in lysis buffer (10 mM Tris-HCl pH 7.4, 10 mM NaCl, 3 mM MgCl₂, 0.1% NP-40) to obtain ∼70,000 individual nuclei. After removing the lysis buffer, a transposition reaction (1.25 μl of Tn5 enzyme in 10 mM Tris-HCl pH 8.0, 5 mM MgCl₂, 10% w/v dimethylformamide) was carried out for 30 min at 37°C, and the resulting fragments were purified using the MinElute PCR Purification Kit (Qiagen). Quantitative PCR was performed to determine the optimal number of amplification cycles for each library, with a unique primer pair assigned to each sample. Libraries were purified using the MinElute PCR Purification Kit (Qiagen), and DNA concentration was measured with an Invitrogen™ Qubit™ 4 Fluorometer using the Qubit 1× dsDNA HS Assay Kit.

### ATAC-seq processing and peak calling

ATAC-seq data from mayfly heads were processed following the pipeline described in (*35*). Briefly, paired-end reads were aligned to the *C. dipterum* reference genome (GCA_902829235.1) using Bowtie2 (*59*). Peak calling was performed with MACS2 (*60*), and Irreproducible Discovery Rate (IDR) analysis (*61*) was used to define condition-specific IDR optimal peak set derived from pooled pseudoreplicates. Consensus peaks from all conditions were merged to create a unified catalogue of accessible putative regulatory elements (APREs), which were annotated to *C. dipterum* genes (clodip_v4 gene models available at (*50*)) using using cisreg_map.py (https://github.com/m-rossello/GeneRegLocator) (Dataset S1). See quality control for these libraries in Fig. S10 and Table S7.

### Gene regulatory network inference using SCENIC

Gene regulatory networks were reconstructed using SCENIC (*33*), implemented in pySCENIC v0.12.1 and run independently per developmental stage. Condition-specific cisTarget databases were generated from ATAC-seq consensus peaks to restrict motif enrichment to accessible regulatory regions. Co-expression modules were inferred with GRNBoost2 (pyscenic grn), tested for motif enrichment against the condition-specific databases (pyscenic ctx), and pruned using custom filtering on enrichment score, motif AUC, orthology identity and target gene count. SCENIC-derived TF–target modules were converted into directed regulatory networks. Target genes were retained as network edges only if they met minimum thresholds for expression level, number of expressing cells, and fraction of expressing cells; thresholds were optimised independently for each cell population to account for differences in cell number and sequencing depth (Dataset S2). Full pipeline details are provided in extended methods.

### Identification of sex-specific regulons

Sex-enriched regulons were identified by applying Fisher’s exact tests to the binarized regulon activity matrix, comparing the proportion of cells with an active regulon between male and female cells. Multiple testing was controlled using Benjamini–Hochberg correction (*62*). Regulons were classified as male- or female-enriched if they met all of the following criteria: adjusted p-value < 0.05 and absolute log₂ fold-change > 2, corresponding to at least a four-fold difference in the proportion of active cells between sexes, where positive and negative values indicate enrichment in males and females respectively.

### Network topology analysis

Gene regulatory networks were analysed to identify three functional roles: hubs, effectors, and initiators. Hub and authority scores (HITS algorithm, (*63*)) were used to classify TFs of disproportionately high connectivity as hubs, and strongly co-regulated genes as effectors. Initiators are a new metric identifying TFs positioned at the entry point of the regulatory cascade, defined as those with no upstream regulator within the network. They are scored by the cumulative hub-weighted influence of their downstream targets. For more information, see extended methods.

### Regulatory network inference using ANANSE

Gene regulatory networks were inferred across embryonic stages of *C. dipterum*. The RNA-seq and ATAC-seq datasets used in this analysis correspond to the embryonic data generated in (*35*) and were processed following the methods described in that study. Network inference was performed with ANANSE (*36*), using its two-step workflow comprising transcription factor binding prediction and regulatory network construction.

### Identification of *DsxM*

We searched the *C. dipterum* genes of interest in the orthogroups previously obtained by (*34*) using OrthoFinder 2 (*64*). The DM domains were identified in eleven insect species using HMMER v3.3.2 with the PFAM DM domain profile (PF00751). The amino acid sequences corresponding to the DM domains were aligned using MAFFT v7.505 with the --localpair and --maxiterate 1000 options. Maximum-likelihood phylogenetic analyses were performed using IQ-TREE v2.0.7 (multicore version) with 1,000 ultrafast bootstrap replicates (-B 1000) and the --mtree option. The best-fitting substitution model was selected using ModelFinder. The phylogenetic reconstruction was carried out with IQ-TREE version 3.0 (*65*). The protein substitution model for each tree was automatically estimated, and node support was calculated from 1000 ultrafast bootstrap replicates. Phylogenetic tree images were obtained using the iTOL webserver (*66*). Trees were rooted using outgroups according to available phylogenetic information.

### Tissue clearing

To visualize the internal soft tissues of *C. dipterum* individuals, we adapted the protocol established in (*67*). Briefly, severed insect heads were fixed in 4% formaldehyde (FA) in phosphate-buffered saline (PBS). After o.n fixation at 4° C, samples were rinsed with PBT (PBS, Triton 0,3%). Heads were bleached in 35% hydrogen-peroxide for 5 days, approximately until pigment was eliminated. After washing the hydrogen-peroxide in PBS, heads were dehydrated in a graded ethanol series (50%, 70%, 90%, 95%, 3 × 100%). Finally, heads were cleared by transferring them into methyl-salicylate (M-2047, Sigma–Aldrich Chemie GmbH, Steinheim, Germany).

### PH3 Immunostaining

Antibody staining on nymphal male heads was done using a standard protocol. Primary antibody used in this study was: rabbit anti-PH3 at 1:1000 (Sigma) and Alexa-Fluor-647 conjugated secondary antibody and rhodamine phalloidin (R415) were from Molecular Probes. Nuclei were detected by 4′,6-diamidino-2-phenylindole (DAPI) staining. Briefly, nymphal heads from male individuals were fixed overnight (o.n.) in 4% FA in PBT [PBS, 0.3% Triton100] at 4 °C. After rinsing the fixative, heads were dissected to remove the external cuticle and to expose the visual system. Incubations with primary and secondary antibodies were performed o.n. at 4 °C. Leica SPE confocal microscope was used to acquire images that were processed with Fiji (*68*).

### *In situ* hybridization

Specific primers were designed to generate DIG-labelled RNA probes against marker genes (Table S14). After overnight (o.n.) fixation of the heads in 4% FA in PBTw [PBS, 0.1% Tween 20] at 4 °C, post-fixated samples were washed with PBTw. In the case of nymphal brains, they were bleached [1.2% H2O2, 0.1X of SSC (3M NaCl, 0.3M Sodium citrate) 5% Formamide] under a bright light overnight and washed thrice with PBTw. Afterwards, they were treated with Detergent Solution (1% SDS, 0.5% Tween, 50 mM Tris-HCl pH 7.5, 1 mM EDTA pH 8 and 150 nM NaCl) for 40 min (embryos) or 2 hours (brains). Then, the samples were washed with PBTw and MABTw [maleic acid buffer (MAB), 0,1% Tween 20] and pre-adapted in hybridisation solution (50% Formamide, 5x SSC, 5% Dextran Sulfate, 0.1% Tween 20, 0.5M EDTA pH 8, 1mg/mL Salmon Sperm DNA, 100 µg/mL heparin) 2h at 60 °C. Afterwards, it was incubated in probe solution (1 ng/μL of probe in hybridisation solution) overnight at 60 °C. After serial washes of decreasing formamide and SSC buffer at 60 °C, the samples were washed twice with MABTw, incubated for 10 minutes with inactivation solution (7.5 mg/mL of glycine, 60 mM HCl, 0.1% Tween 20) and washed thrice with MABTw again. Then, they were pre-adapted in blocking [1% Blocking reagent (Roche) in MABTw] and incubated in antibody solution [1/2000 Anti-DIG-AP (Sigma-Aldrich) in Blocking solution] overnight at 4 °C. Then, the samples were washed 4 times with MABTw, pre-adapted twice with AP buffer [100mM Tris pH 9.5, 50mM MgCl2, 100mM NaCl, 0.1% Tween 20] and incubated in development solution [100mM Tris pH 9.5, 50mM MgCl2, 100mM NaCl, 0.1% Tween 20, 6.9% Polyvinyl alcohol, 10 µl/ml NBT/BCIP (Sigma-Aldrich)] until the signal was clear. Then, the reaction was stopped with PBS, samples were post-fixed in 4% FA in PBS for 20 min and the background signal was removed by a fast wash in Ethanol. Afterwards, the sample was mounted on glycerol and the images we were taken either in a Leica M205 FCA stereoscope (brains) or a Nikon Eclipse Ci optical microscope (embryos) and edited using Fiji (*68*).

### HCR *in situ* hybridization

HCR hybridization followed a modified version of the Molecular Instruments HCR v.3 protocol (https://hackmd.io/@ColbyMBL/hcr (*69*)). The HCR probe was designed to evade non-specific binding using an open-source probe design program (*70*). Briefly, samples stored in ethanol were rehydrated in an ethanol series (75%, 50%, 25%) in PBTw. After 3× 5 min washes in PBTw, samples were permeabilized in detergent solution for 40 min at room temperature (RT), kept in pre-warmed Probe Hybridization Buffer (Molecular Instruments) for 30 min at 37°C, and incubated in Probe Solution [12 nM of probe in Probe Hybridization Buffer (Molecular Instruments)] overnight at 37°C. After 4×15 min washes in pre-heated wash buffer (Molecular Instruments) at 37°C and 2×5 min washes in 5× SSCTw (SSC, 0.1% Tween 20) at RT, they were kept in pre-equilibrated Amplification Buffer (Molecular Instruments) for 30 min at RT and incubated overnight in the dark at RT in hairpin solution [60 nM of each hairpin h1 and h2 (Molecular Instruments) separately in pre-equilibrated Amplification Buffer], that was previously heated at 95°C for 90 s and cooled down for 30 min. Following 5×20 min washes in 5× SSCTw and 1×10 min wash in PBTw 0.1% pH 7.4 in the dark at RT, samples were incubated for 1h at RT or overnight at 4°C in PBTw 0.1% pH 7.4 with DAPI (1 µg/mL) and mounted on glycerol. Images were acquired using a Zeiss LSM 880 confocal microscope and were processed with Fiji (*68*).

### RNAi analysis and genotyping

RNAi synthesis. RNAi was synthetised as described in (*71*), with several modifications. gfp and dsxM constructs had the T7 (5’-TAATACGACTCACTATAGGG-3’) and SP6 (5’-ATTTAGGTGACACTATAGA-3’) promoters. The primers used for amplification were the M13F (5’-GTAAAACGACGGCCAG-3’) and M13R (5’-CAGGAAACAGCTATGAC-3’).

Females were fertilized as previously described in (*38*). Six days after fertilization, females were injected through the joint between the 6th and the 7th abdominal segment twice, one injection per egg sack. Then, they were kept at stable conditions for 2 days post injection (dpi) and the embryos were collected by dissecting the gravid females.

After a colorimetric *in situ* hybridisation, each embryo was collected separately and transferred in 50 µL of QuickExtrac^TM^ DNA Extraction Solution 1.0 (Biosearch technologies). Then, the samples were vortexed for 15s, and incubated at 65°C for 15 minuts, 68°C for 15 minutes and 98°C for 10 minutes. Male and female individuals were identified by PCR using the dsxM_F3 primer (CCTCGCCCCTTGAACCAGAGGA) and the dsxM_R3 primer (CACAAAACCAAGCGCAGGCACC), as only male showed an amplified region. For control, a region of a *yellow* gene was amplified using y_F1 (AAATCGTGTTTCCGCACAGG) and y_R1 (AGCGAACGTTTCCTTCAACA) primers.

### DIrect-PArental CRISPR/Cas9 (DIPA-CRISPR) targeted mutagenesis

We used the CRISPOR website (https://crispor.gi.ucsc.edu/) to design crRNA from the *C. dipterum* genome (GCA_902829235.1) (*72*). A specific guide (TTCAAGATACACGCAATATC) targeting the DNA binding domain of *dsxM* was selected after assessing the absence of off-targets by BLAST against the genome. Selected crRNA (2 nmol, IDT) and tracrRNA (#1072533, IDT) were resuspended at 200 µM. 1 µL of each were annealed together at 95°C for 5 minutes to obtain a pgRNA (crRNA::tracrRNA) at 100µM. *C. dipterum* females crossed and immediately injected (within an hour) as described above using the following solution: pgRNA_01 (22 µM), Cas9 protein (18,3µM, #1081058, IDT), Hepes (10 mM), KCl (75 mM), Triton (1%) and Dextran-Fluorescein (#D1821, ThermoFisher).

For control individuals, a pgRNA against a yellow gene (TTCCACGCGTTCGACCCTCG) having no off target on dsxM locus was used as replacement with the same injection solution described above.

Injected females were kept at stable conditions and nymphs were collected after hatchings and maintained in an isolated beaker (as described above).

### Genotyping and mutant screening

As DIPA-CRISPR is an indirect injection method, several individuals were isolated on (24-well plates) and genomic DNA was extracted from one leg of each individual (at nymphal NIII stage) in 20µL of QuickExtract™ DNA Extraction Solution (Biosearch technologies) according to the provider’s protocol. Male and female individuals were identified by PCR using the dsxM_F3 primer and the dsxM_R3 primer, as only male showed an amplified region. Mutations in males were screened by Sanger sequencing using the dsxM_F3 primer. Males showing possible mutations were isolated and their development monitored to assess presence of phenotypes during their development. Images of individuals were taken using the Nikon SZM-745 stereoscope for nymphs or the Leica M205-FCA stereoscope for adults and edited using Fiji (*68*).

**Fig. S1. Quality control and cluster composition of the SPLiT-seq dataset. (A)** UMAP embedding of cells for each condition, with replicates shown in different shades of the same colour. **(B-C)** Violin plots showing the number of genes detected (B) and total UMI counts (C) per cell for each replicate, grouped by condition. **(D)** Proportion of cells from each replicate contributing to each cluster. **(E-F)** Violin plots showing the number of genes detected (E) and total UMI counts (F) per cell for each cluster. **(G)** Proportion of cells from each cluster contributing to each replicate, grouped by condition.

**Fig. S2.** UMAP of gene markers per cluster. Marker gene expression projected onto the UMAP, min-max scaled to [0, 10].

**Fig. S3.** Quality control and cluster composition of the visual system sub-clustering. **(A)** UMAP embedding of cells for each condition, with replicates shown in different shades of the same colour. **(B-C)** Violin plots showing the number of genes detected (B) and total UMI counts (C) per cell for each replicate, grouped by condition. **(D)** Proportion of cells from each replicate contributing to each cluster. **(E)** UMAP embedding of the eye lineage subclusters, coloured by cell population (optic lobe, developing retina, photoreceptor), with arrows indicating PAGA connectivity between clusters. **(F-G)** Violin plots showing the number of genes detected (F) and total UMI counts (G) per cell for each cluster. **(H)** Proportion of cells from each cluster contributing to each replicate, grouped by condition.

**Fig. S4.** Spatial expression of visual system cell clusters. Expression of main markers of visual system clusters detected through in situ hybridization in male and female nymphal heads.

**Fig. S5.** Expression of the mayfly opsins in the visual system clusters. **(A)** Dot plot of expression, by mean normalised counts, of all described mayfly opsins across all visual system clusters. **(B)** UMAP representation of opsin expression; only opsins with detectable expression are shown.

**Fig. S6.** Proportion of cells in each cell cycle phase. Proportion of cells predicted to be in G1, S, and G2M phases shown as individual panels for each combination of cell cycle phase and nymphal stage, grouped by cell population (A) or cluster (B). Error bars represent the standard deviation across biological replicates.

**Fig. S7.** Cell neighbourhood enrichment analysis. **(A-B)** Neighbourhoods with spatialFDR < 0.05 (A) and spatialFDR < 0.01 (B). Teal neighbourhoods represent significantly male-enriched neighbourhoods and purple ones female-enriched. **(C-D)** Violin plot representations of all neighbourhoods by log fold change, grouped by cell population (C) or cluster (D). **(E-F)** Volcano plots of neighbourhood enrichment, grouped by cell population (E) or cluster (F). In violin plots, black dots represent spatialFDR > 0.01 and grey dots spatialFDR < 0.05. Neighbourhood enrichment analysis performed using Milo. The “Mixed” group comprises neighbourhoods that cannot be associated with any single cell population or cluster, as they contain cells from multiple groups.

**Fig. S8.** Volcano plot representation of the pseudo-bulk differential gene expression (pDGE). Each panel shows log fold change against -log10(P value) for one cell population at Nymph V (left) and Nymph VII (right), with dashed lines marking the significance thresholds (P < 0.05, |logFC| ≥ 1.5). Genes passing these thresholds are coloured in teal (male-biased) or purple (female-biased), with significant transcription factors highlighted as larger, labelled dots.

**Fig. S9.** Identification of dsxM as a *C. dipterum*-specific dsx paralogue. (A) Multiple sequence alignment of the DM domain of dsx family proteins across insect species. **(B)** Maximum likelihood phylogenetic tree with bootstrap support values, showing three dsx paralogues in *C. dipterum* (*dsx*, *dsxM*, and *dsxl*, highlighted in bold), with dsxM forming a distinct clade. **(C)** Agarose gel electrophoresis of PCR products from genomic DNA of males and females, using yellow as a positive control (C+) and a no-template reaction as negative control (C−).

**Fig. S10. Fragment size distribution of the ATAC-seq libraries.** Normalised read density as a function of fragment length (bp) for each biological replicate across conditions. Insets show the same data on a logarithmic scale.

**Fig. S11.** Accessible Putative Regulatory Elements (APREs) identified by ATAC-seq visualised in the *C. dipterum* genome browser. Example locus showing the so gene region. The first track shows consensus APREs (black bars) identified from IDR-filtered peaks across all conditions. Subsequent tracks show normalised ATAC-seq signal for each condition.

**Fig. S12.** Enriched sex-specific regulons in the visual system. For each tissue and stage, bar plots show female-enriched (left) and male-enriched (right) regulons, ranked by percentage of cells with an active regulon, as identified by Fisher’s exact test. The scatter plot (centre) shows regulon prevalence by sex, coloured by female-biased (purple), male-biased (teal), or not significant (grey).

**Fig. S13.** Network topology analysis of gene regulatory networks in the different cell populations of the visual system. **(A)** Initiator score scatter plots for all regulons. **(B)** Hub score scatter plots for all regulons. **(C)** Authority score scatter plots for all target genes. Plots are shown for the developing retina and optic lobe at Nymph V and Nymph VII. Coloured dots represent nodes passing the threshold for each category, with the highest-scoring nodes labelled.

**Fig. S14.** ANANSE edge score distributions. Filtering probability threshold retains high-confidence TF–target interactions while preserving the overall score distributions. Values before (left) and after (right) probability threshold filtering. Density plots show the distribution of the predicted regulatory probability (prob), TF binding score (weighted_binding), TF activity (activity) and gene expression (target_expression) for all TF–target edges.

**Fig. S15**. **Regulation of the RDGN by dsxM during embryogenesis.** Network graphs showing the regulatory relationships between dsxM and RDGN genes across embryonic stages 4 to 14, inferred from whole-embryo bulk RNA-seq and bulk ATAC-seq data using ANANSE. Thicker edges represent direct connections from an initiator to an RDGN gene.

**Fig. S16.** Quantification of *ey* area following dsxM knockdown by dsRNA injection. Representative brightfield images of *ey* expression domain in dsRNA-gfp (control) and dsRNA-dsxM individuals, with measured *ey* expression domain outlined (top). *ey* expression domain quantification for all samples, in pixels (bottom). Rows in bold and marked with an asterisk correspond to the individuals shown above.

**Fig. S17.** DIPA-CRISPR mutagenesis of dsxM and yellow. Eye phenotype (arrowhead), sgRNA target locus, and Sanger sequencing chromatogram of the mutated sequence for a dsxM mosaic F0 individual (top). Eye phenotype, hatchling phenotype showing loss of pigmentation (arrowhead), sgRNA target locus, and Sanger sequencing chromatogram for a yellow mosaic F0 individual (bottom).

**Fig. S18.** Graphical representation of the modified SCENIC pipeline used in this study. This pipeline combines scRNA-seq with bulk ATAC-seq to identify active regulons in each cell and quantify their activity. Co-expression modules are inferred between transcription factors (TF) and genes across cells; motif discovery refines regulons by retaining targets with supporting motif evidence; cell scoring computes an AUC per cell based on the expression rank of each regulon’s target genes; binarisation converts continuous AUC scores into active/inactive regulon calls across cells.

## List of Tables

**Table S1**. BarQC statistics. scRNA-seq statistics

**Table S2.** Cluster markers. Top 5 markers for each cell cluster

**Table S3.** Cluster annotation. Cluster annotations based on top markers.

**Table S4.** Eye Cluster markers. Top 5 markers for each cell cluster from the visual system.

**Table S5.** pseudobulk-DGE expression in the different cell types **Table S6.** pseudobulk-DGE expression in the visual system cell types. **Table S7.** ATAC-seq peak calling statistics

**Table S8.** Visual system regulons

**Table S9.** Visual system regulons enriched

**Table S10.** Topological metrics of the male regulons and targets

**Table S11**. GO term enrichment of male effectors. **Table S12.** Dsx binding motifs in *eyeless* locus **Table S13.** Decision table for TF curation.

**Table S14**. In situ probes.

## List of Datasets

**Data S1.** (separate file). Consensus accessible putative regulatory elements (APREs) identified by ATAC-seq. One BED file per condition (NIII, NV female, NV male, NVII female, NVII male) containing IDR-filtered consensus peaks.

**Data S2.** (separate file). SCENIC network data. Edge and node tables of the final filtered SCENIC output. For each condition, files are provided separately for the developing retina and optic lobe tissues.

**Data S3.** (separate file). ANANSE filtered network data. Edge and node tables of the probability threshold-filtered TF–target interactions.

## Notes

### Competing Interest Statement

The authors have declared no competing interest.

