## Supplementary information and figures for "A sex-specific regulator expands the embryonic ocular field to generate a novel visual system"

**Suppl Tables and Figures**

**Tables**

**Table S1. BarQC statistics. scRNA-seq statistics**

**Table S2. Cluster markers. Top 5 markers for each cell cluster**

**Table S3. Cluster annotation. Cluster annotations based on top markers.**

**Table S4. Eye Cluster markers. Top 5 markers for each cell cluster from the visual system.**

**Table S5. pseudobulk-DGE expression in the different cell types**

**Table S6. pseudobulk-DGE expression in the visual system cell types.**

**Table S7. ATAC-seq peak calling statistics**

**Table S8. Visual system regulons**

**Table S9. Visual system regulons enriched**

**Table S10. Topological metrics of the male regulons and targets**

**Table S11. GO term enrichment of male effectors.**

**Table S12. *Dsx* binding motifs in *ey* locus**

**Table S13. In situ probes.**

**Table S14. Decision table for TF curation.**

**Supplementary Figures**

**Fig. S1. Quality control and cluster composition of the SPLiT-seq dataset.** (A) UMAP embedding of cells for each condition, with replicates shown in different shades of the same colour. (B-C) Violin plots showing the number of genes detected (B) and total UMI counts (C) per cell for each replicate, grouped by condition. (D) Proportion of cells from each replicate contributing to each cluster. (E-F) Violin plots showing the number of genes detected (E) and total UMI counts (F) per cell for each cluster. (G) Proportion of cells from each cluster contributing to each replicate, grouped by condition.

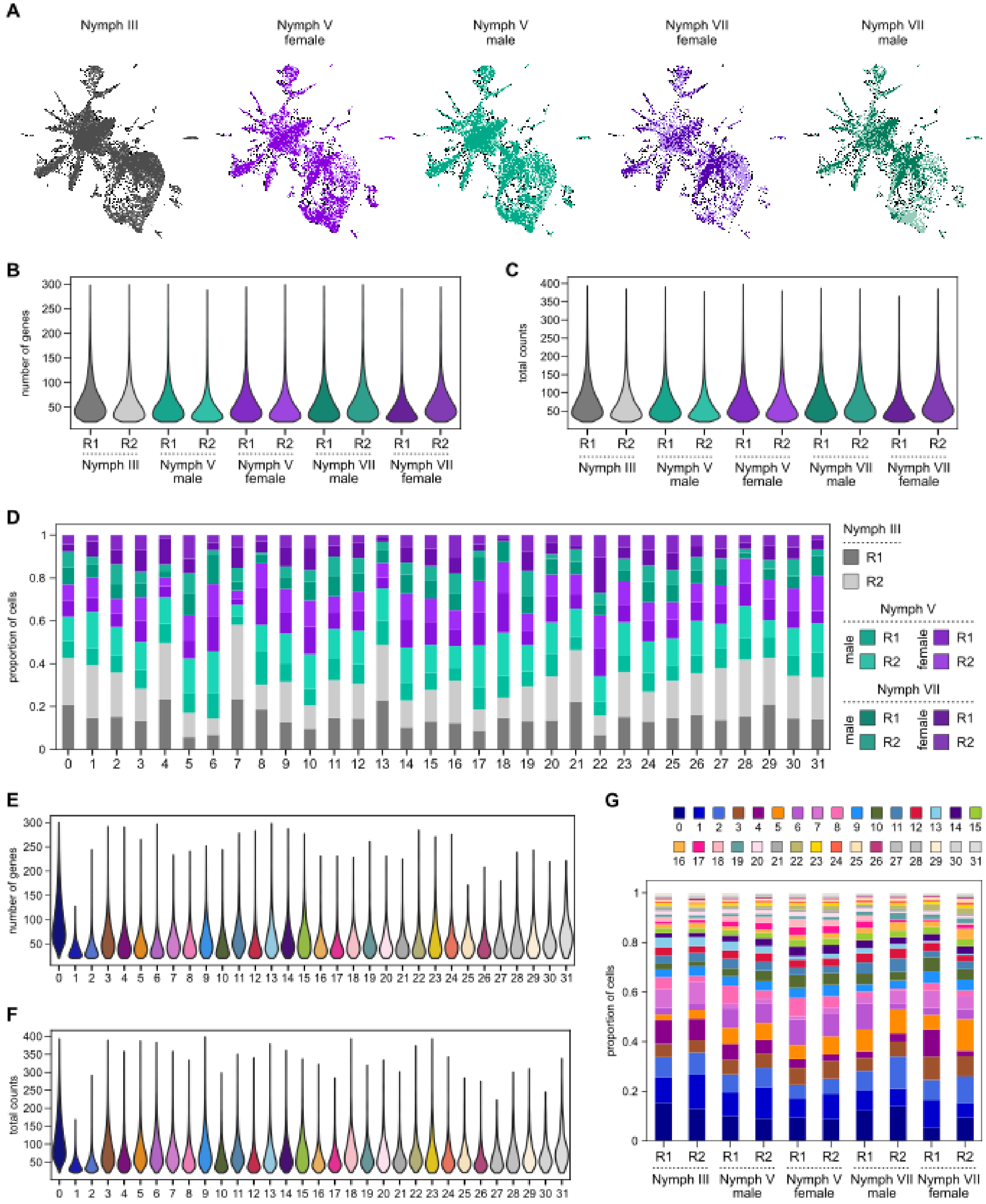

**Fig. S2.** UMAP of gene markers per cluster. Marker gene expression projected onto the UMAP, min-max scaled to [0, 10].

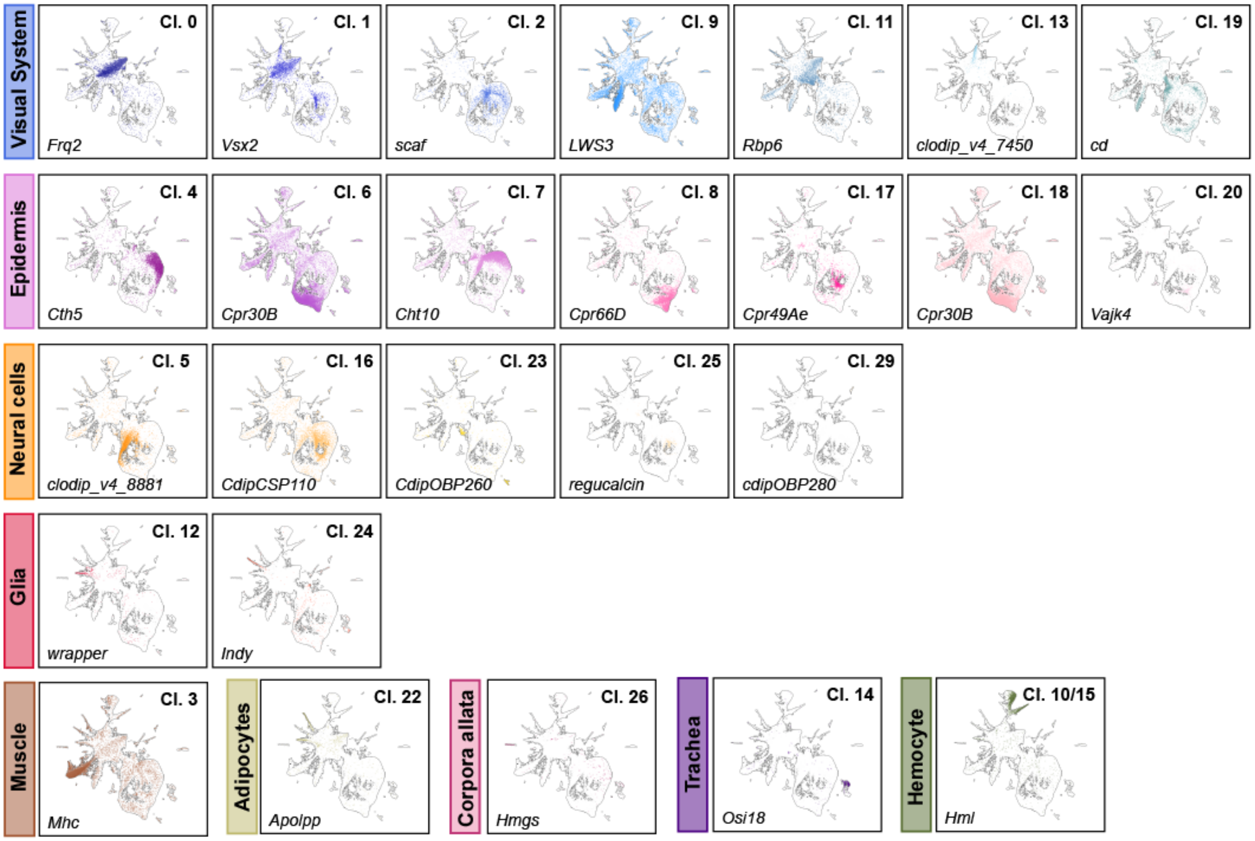

**Fig. S3. Quality control and cluster composition of the visual system sub-clustering.** (A) UMAP embedding of cells for each condition, with replicates shown in different shades of the same colour. (B-C) Violin plots showing the number of genes detected (B) and total UMI counts (C) per cell for each replicate, grouped by condition. (D) Proportion of cells from each replicate contributing to each cluster. (E) UMAP embedding of the eye lineage subclusters, coloured by cell population (optic lobe, developing retina, photoreceptor), with arrows indicating PAGA connectivity between clusters. (F-G) Violin plots showing the number of genes detected (F) and total UMI counts (G) per cell for each cluster. (H) Proportion of cells from each cluster contributing to each replicate, grouped by condition.

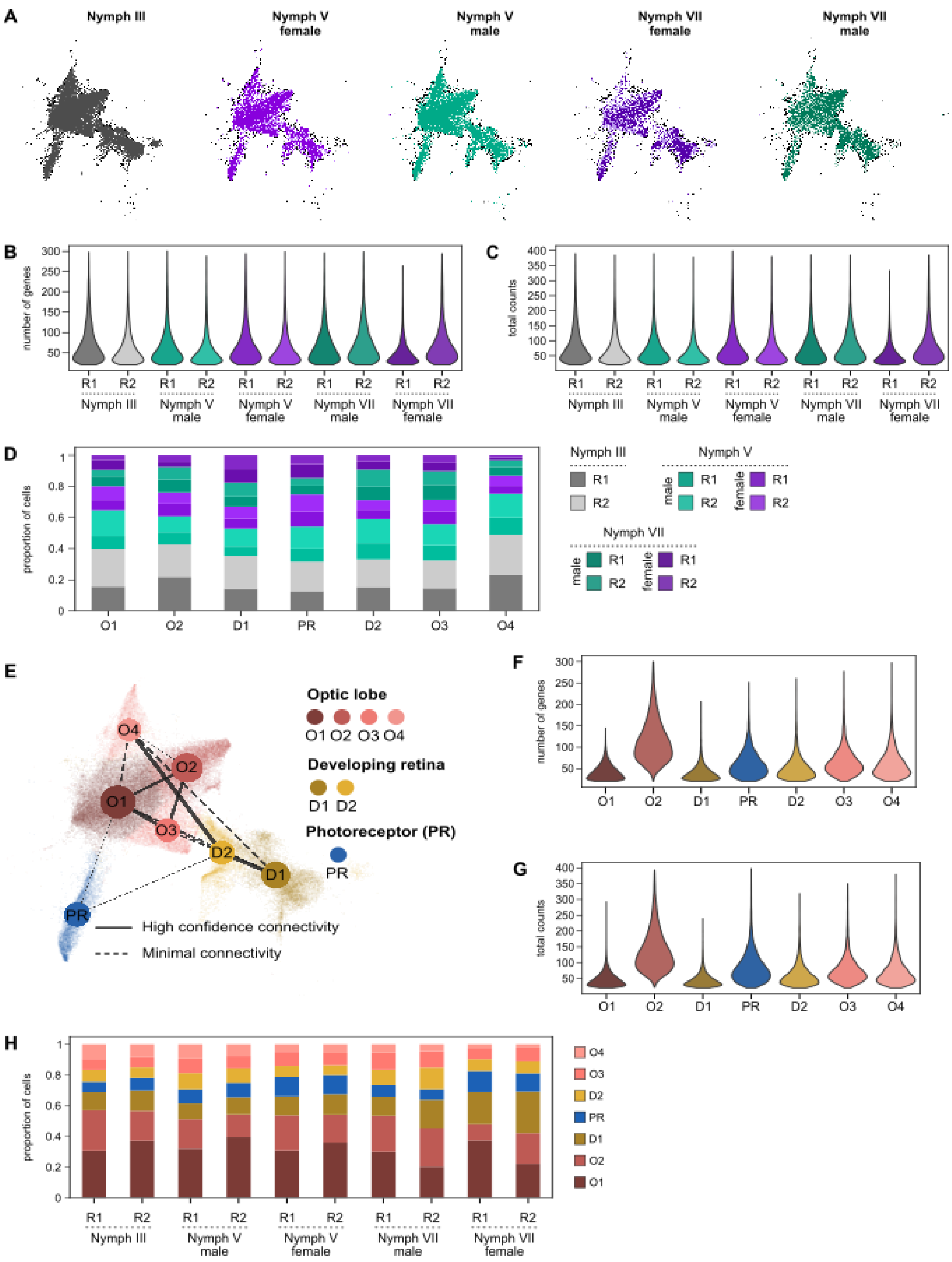

**Fig. S4. Expression of main markers of visual system clusters.** In situ hybridization in male and female nymphal optic systems.

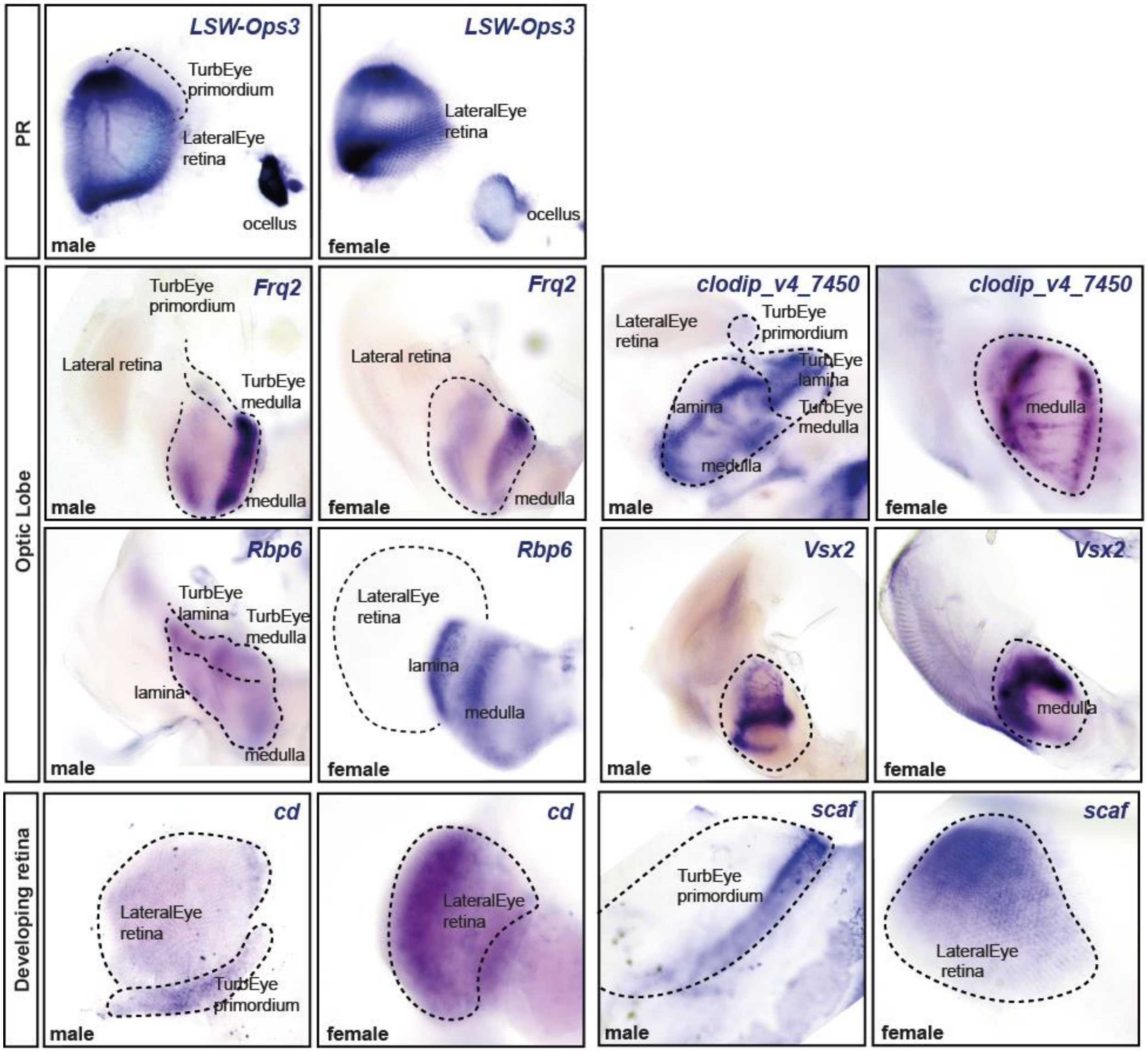

**Fig. S5. Expression of the mayfly opsins in the visual system clusters.** (A) Dot plot of expression, by mean normalised counts, of all described mayfly opsins across all visual system clusters. (B) UMAP representation of opsin expression; only opsins with detectable expression are shown.

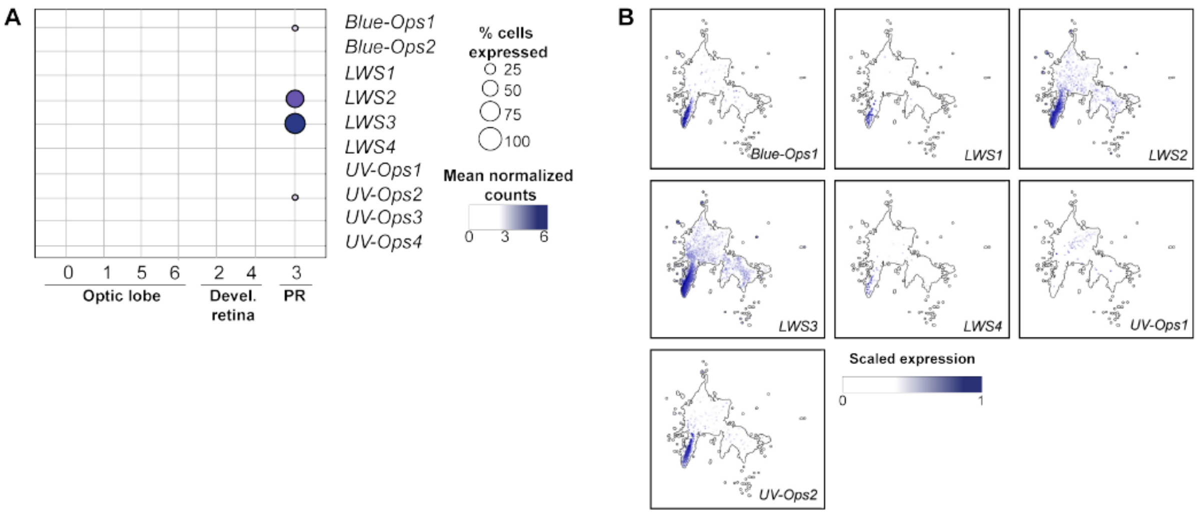

**Fig. S6. Proportion of cells in each cell cycle phase.** Proportion of cells predicted to be in G1, S, and G2M phases shown as individual panels for each combination of cell cycle phase and nymphal stage, grouped by cell population (A) or cluster (B). Error bars represent the standard deviation across biological replicates.

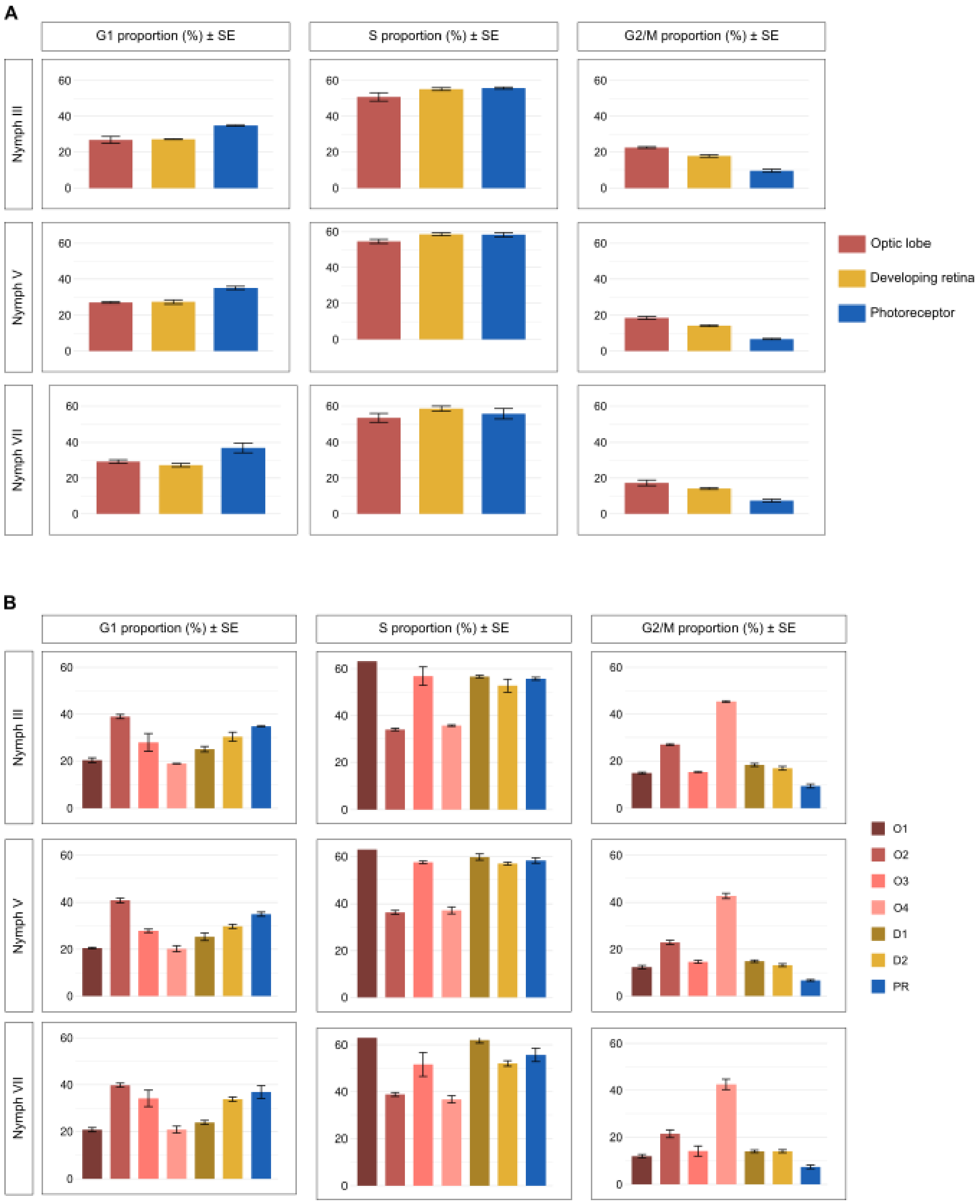

**Fig. S7. Cell neighbourhood enrichment analysis.** (A-B) Neighbourhoods with spatialFDR < 0.05 (A) and spatialFDR < 0.01 (B). Teal neighbourhoods represent significantly male-enriched neighbourhoods and purple ones female-enriched. (C-D) Violin plot representations of all neighbourhoods by log fold change, grouped by cell population (C) or cluster (D). (E-F) Volcano plots of neighbourhood enrichment, grouped by cell population (E) or cluster (F). In violin plots, black dots represent spatialFDR > 0.01 and grey dots spatialFDR < 0.05. Neighbourhood enrichment analysis performed using Milo. The "Mixed" group comprises neighbourhoods that cannot be associated with any single cell population or cluster, as they contain cells from multiple groups.

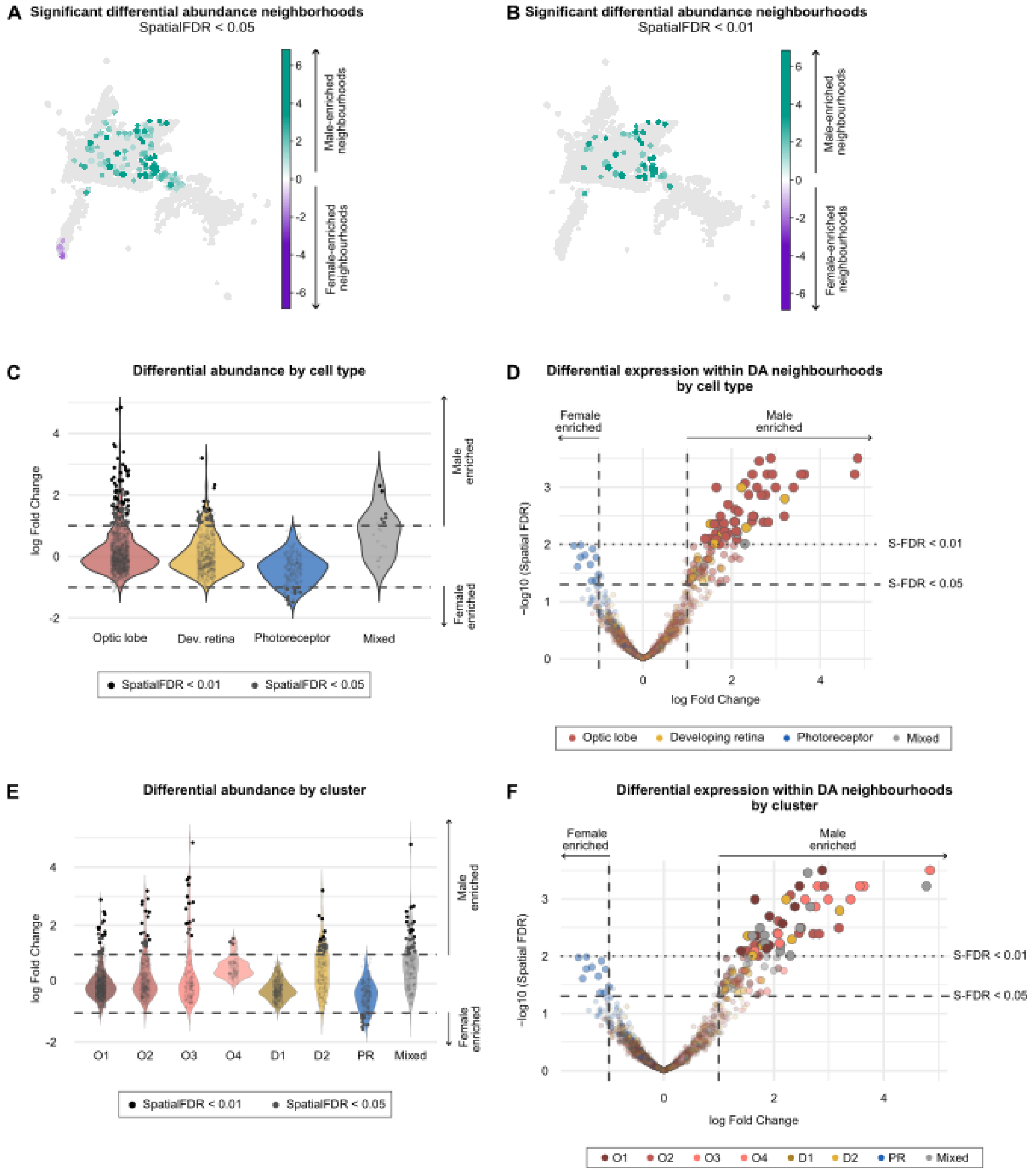

**Fig. S8 Volcano plot representation of the pseudo-bulk differential gene expression (pDGE).** Each panel shows log fold change against -log10(P value) for one cell population at Nymph V (left) and Nymph VII (right), with dashed lines marking the significance thresholds (P < 0.05, |logFC| ≥ 1.5). Genes passing these thresholds are coloured in teal (male-biased) or purple (female-biased), with significant transcription factors highlighted as larger, labelled dots.

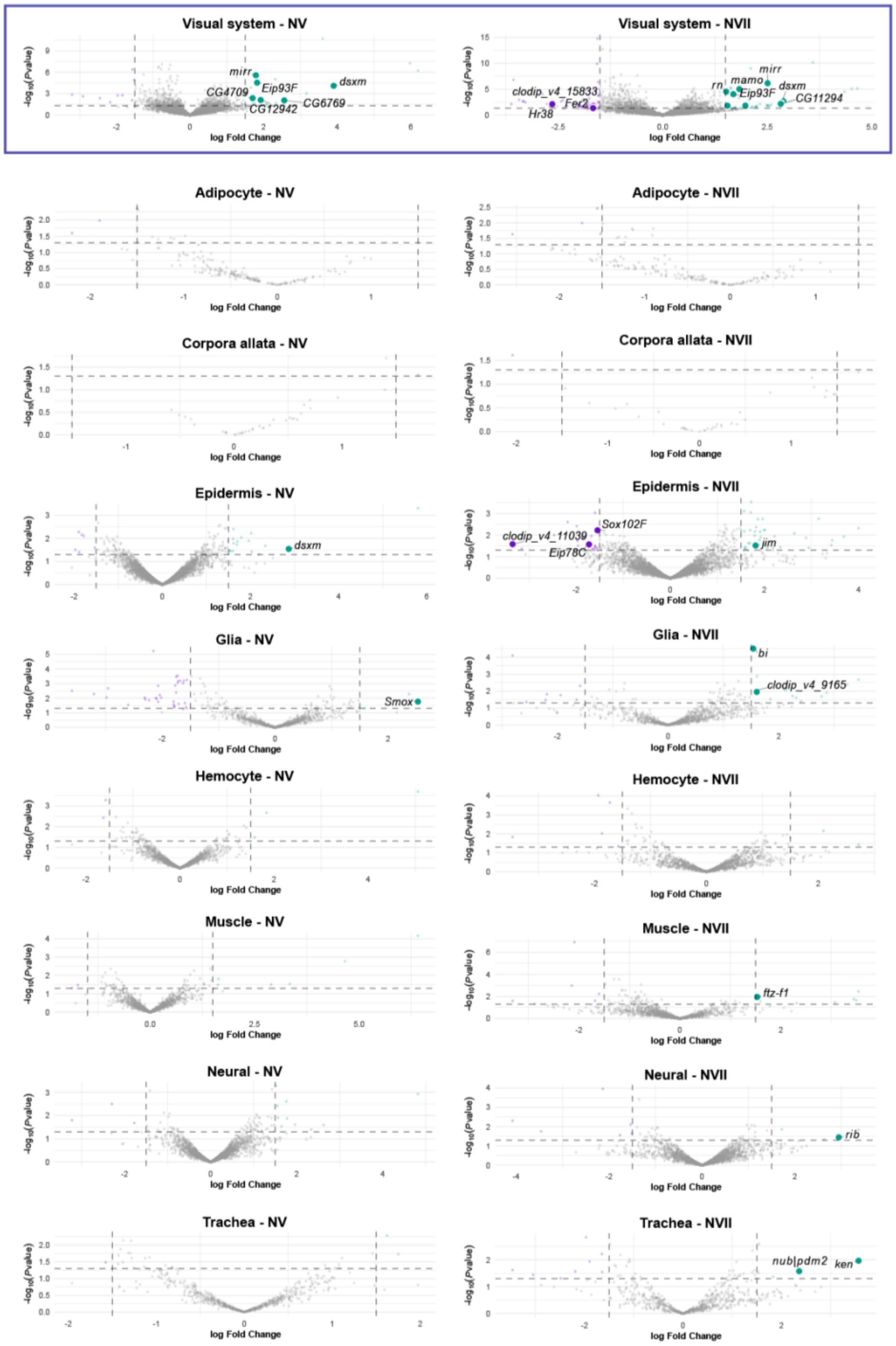

**Fig. S9. Identification of dsxM as a *C. dipterum*-specific dsx paralogue.** (A) Multiple sequence alignment of the DM domain of dsx family proteins across insect species. (B) Maximum likelihood phylogenetic tree with bootstrap support values, showing three dsx paralogues in *C. dipterum* (*dsx*, *dsxM*, and *dsxl*, highlighted in bold), with dsxM forming a distinct clade. (C) Agarose gel electrophoresis of PCR products from genomic DNA of males and females, using yellow as a positive control (C+) and a no-template reaction as negative control (C−).

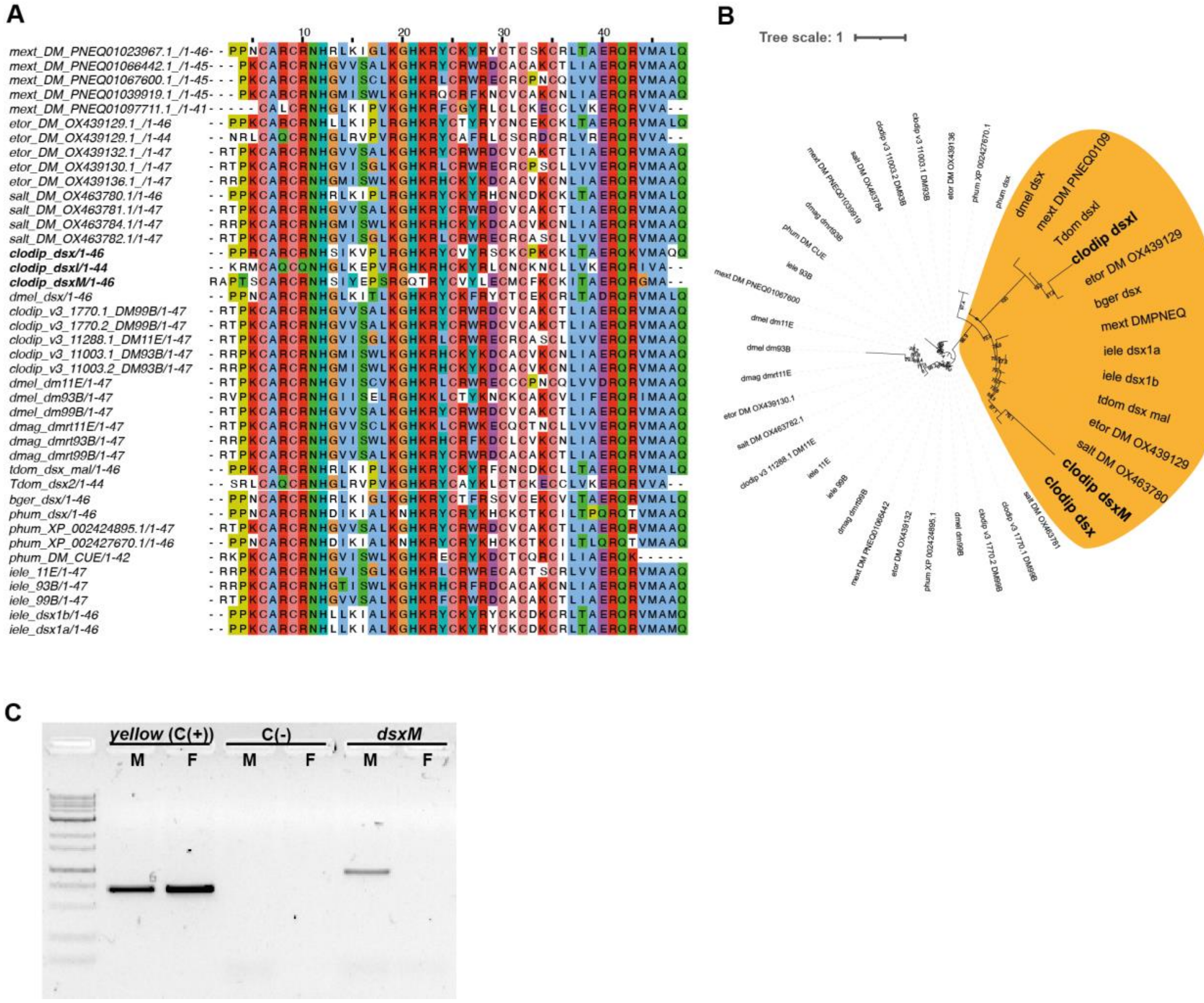

**Fig. S10. Fragment size distribution of the ATAC-seq libraries.** Normalised read density as a function of fragment length (bp) for each biological replicate across conditions. Insets show the same data on a logarithmic scale.

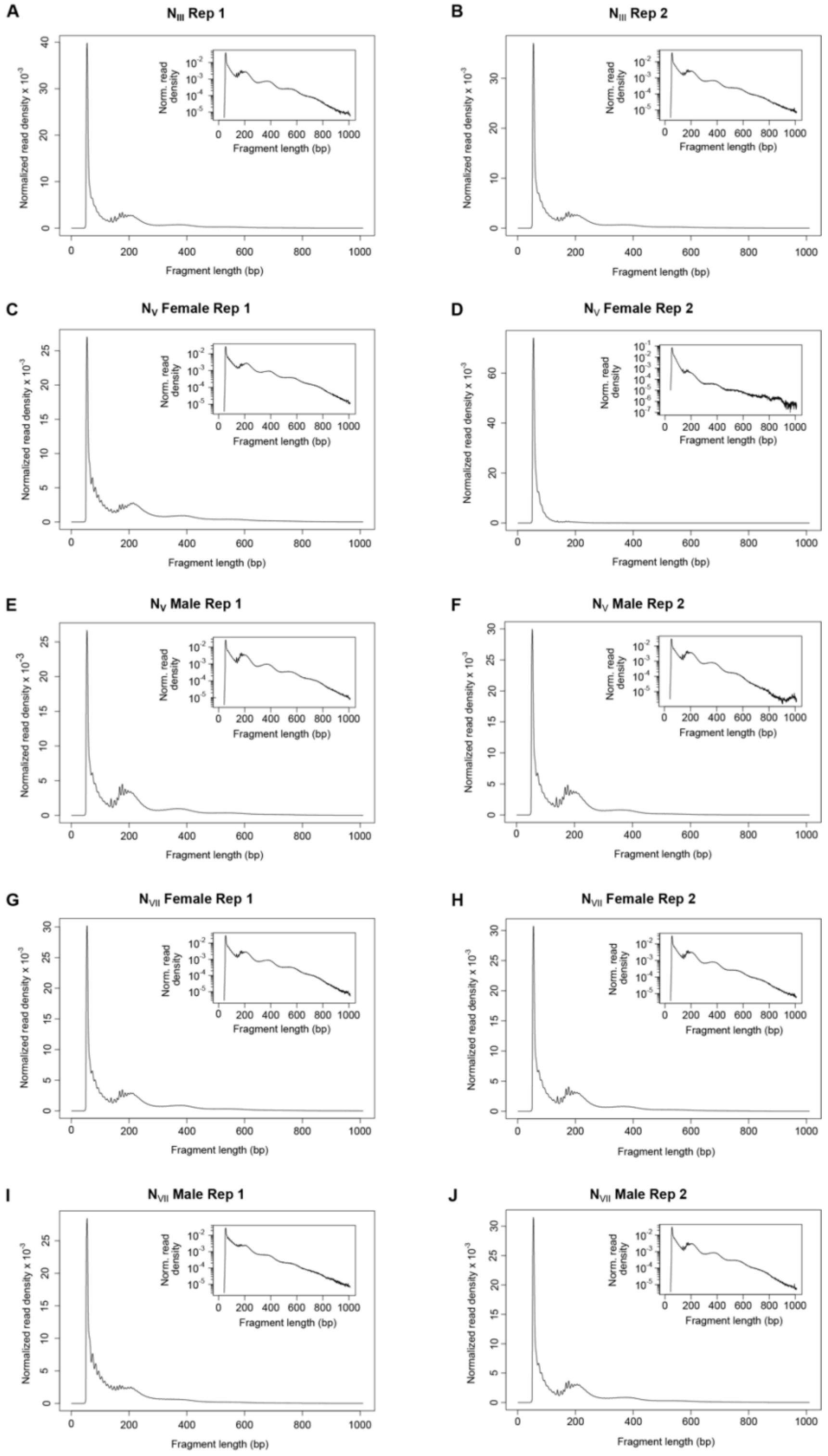

**Fig. S11.** Accessible Putative Regulatory Elements (APREs) identified by ATAC-seq visualised in the *C. dipterum* genome browser. Example locus showing the so gene region. The first track shows consensus APREs (black bars) identified from IDR-filtered peaks across all conditions. Subsequent tracks show normalised ATAC-seq signal for each condition.

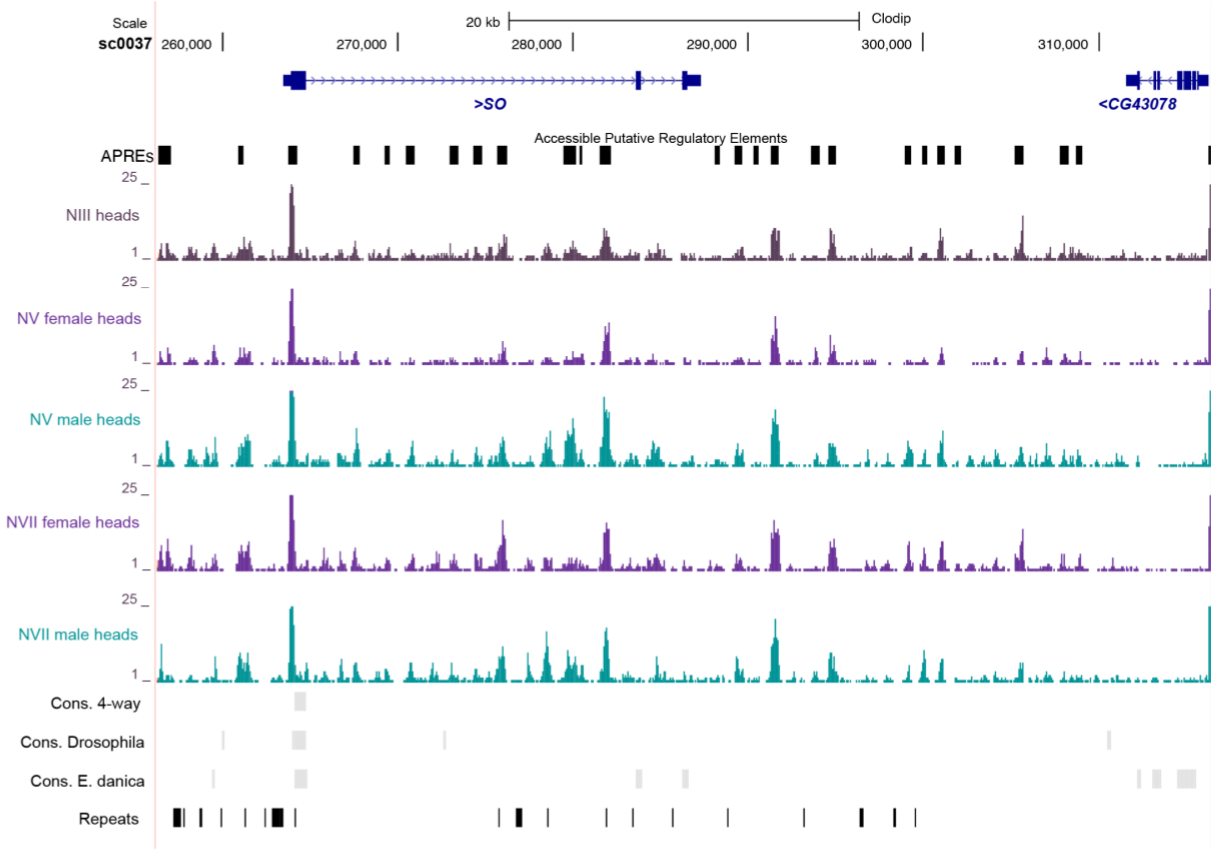

**Fig. S12. Enriched sex-specific regulons in the visual system.** For each tissue and stage, bar plots show female-enriched (left) and male-enriched (right) regulons, ranked by percentage of cells with an active regulon, as identified by Fisher's exact test. The scatter plot (centre) shows regulon prevalence by sex, coloured by female-biased (purple), male-biased (teal), or not significant (grey).

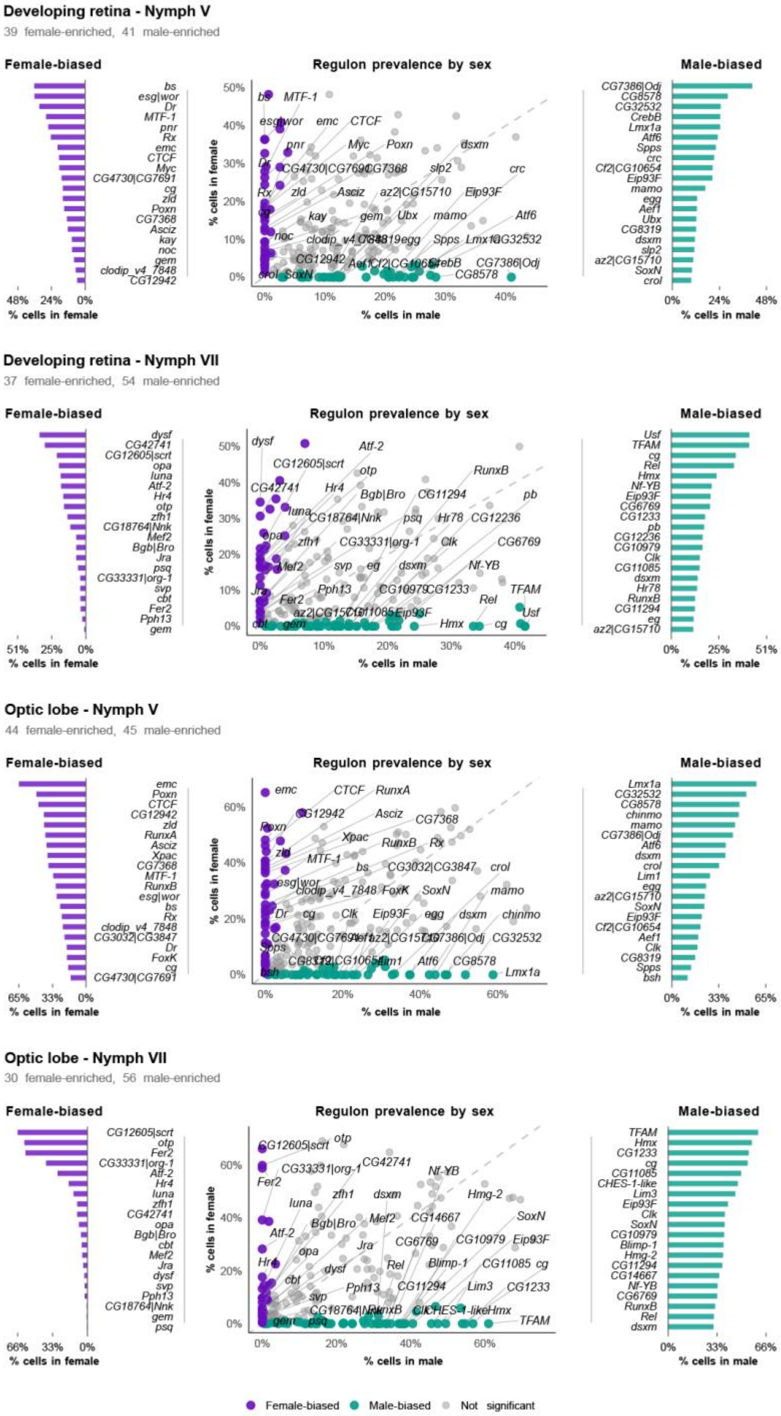

**Fig. S13. Network topology analysis of gene regulatory networks in the different cell populations of the visual system.** (A) Initiator score scatter plots for all regulons. (B) Hub score scatter plots for all regulons. (C) Authority score scatter plots for all target genes. Plots are shown for the developing retina and optic lobe at Nymph V and Nymph VII. Coloured dots represent nodes passing the threshold for each category, with the highest-scoring nodes labelled.

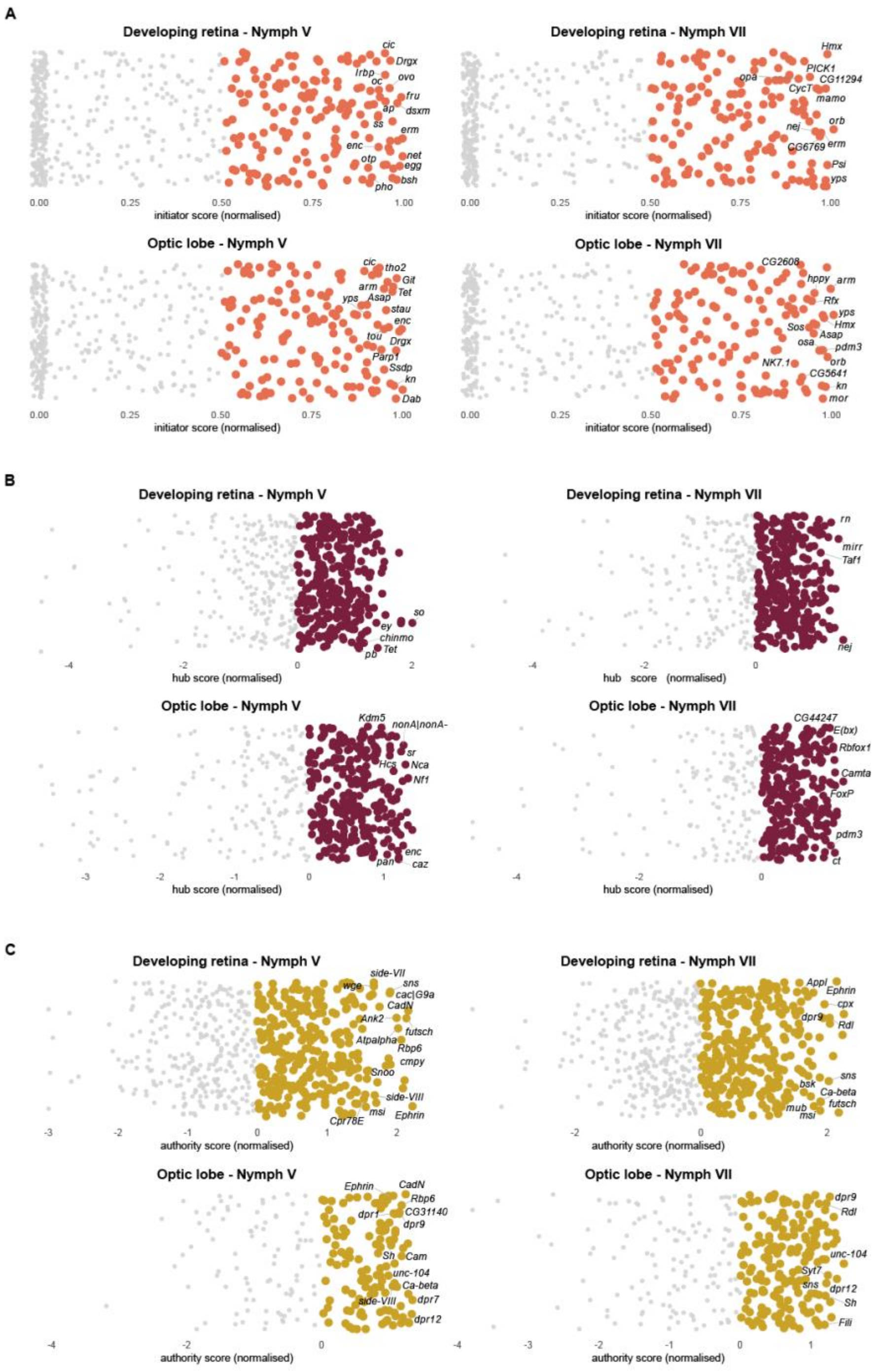

**Fig. S14. ANANSE edge score distributions.** Filtering probability threshold retains high-confidence TF–target interactions while preserving the overall score distributions. Values before (left) and after (right) probability threshold filtering. Density plots show the distribution of the predicted regulatory probability (prob), TF binding score (weighted_binding), TF activity (activity) and gene expression (target_expression) for all TF–target edges.

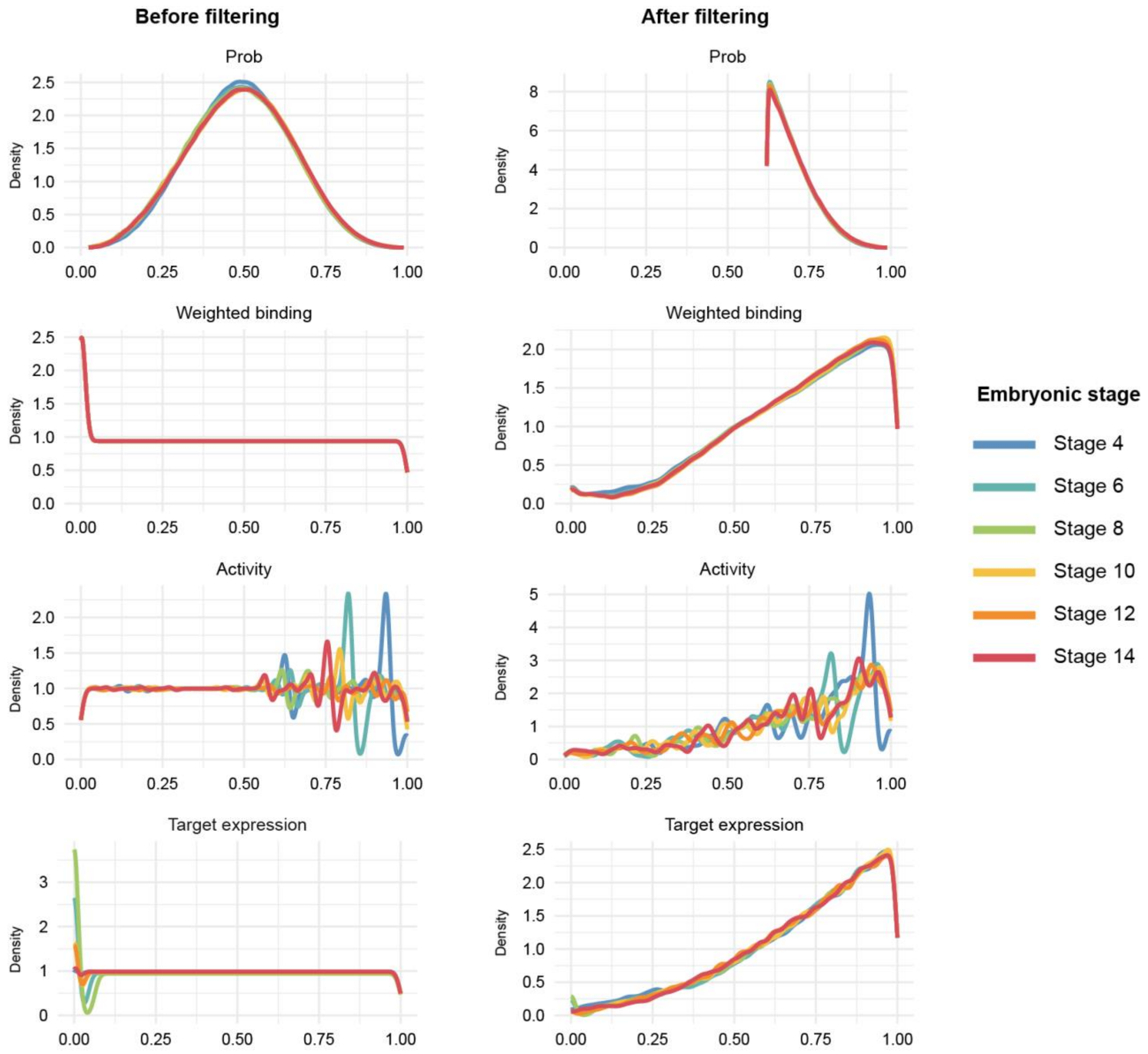

**Fig. S15. Regulation of the RDGN by dsxM during embryogenesis.** Network graphs showing the regulatory relationships between dsxM and RDGN genes across embryonic stages 4 to 14, inferred from whole-embryo bulk RNA-seq and bulk ATAC-seq data using ANANSE. Thicker edges represent direct connections from an initiator to an RDGN gene.

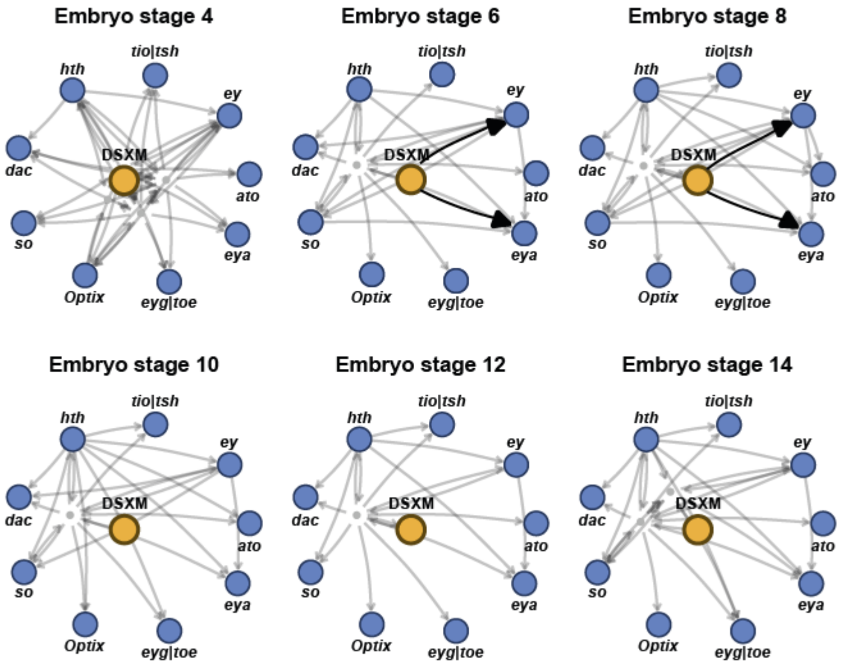

**Fig. S16. Quantification of *ey* area following dsxM knockdown by dsRNA injection.** Representative brightfield images of *ey* expression domain in dsRNA-*gfp* (control) and dsRNA-*dsxM* individuals, with measured *ey* expression domain outlined (top). *ey* expression domain quantification for all samples, in pixels (bottom). Rows in bold and marked with an asterisk correspond to the individuals shown above.

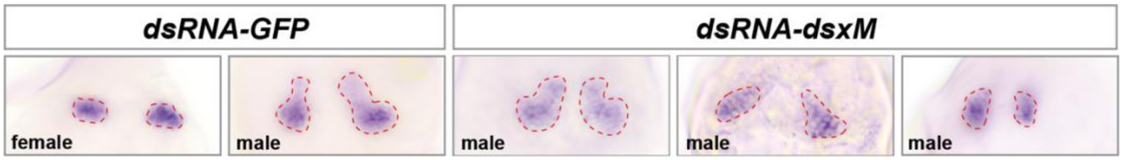


| Sample | Area (px) |
|---|---|
| female GFP | 5401 |
| female GFP | 4628 |
| **female GFP*** | **8257** |
| **female GFP*** | **8133** |
| female GFP | 11064 |
| female GFP | 8975 |
| female GFP | 7763 |
| male GFP | 20721 |
| **male GFP*** | **20049** |
| **male GFP*** | **15059** |
| male_GFP | 19995 |
| male_GFP | 20762 |
| male GFP | 11839 |
| male GFP | 9803 |
| male GFP | 12067 |
| male GFP | 16453 |
| male GFP | 14520 |

| Sample | Area (px) |
|---|---|
| male dsxM | 15205 |
| male dsxM | 11781 |
| male dsxM | 9401 |
| male dsxM | 11430 |
| male dsxM | 13782 |
| male dsxM | 15799 |
| male dsxM | 12046 |
| male dsxM | 9386 |
| male dsxM | 13862 |
| male dsxM | 14063 |
| **male dsxM*** | **13678** |
| **male dsxM*** | **14551** |
| **male dsxM*** | **12006** |
| **male dsxM*** | **12034** |
| male dsxM | 5625 |
| male dsxM | 7543 |
| **male dsxM*** | **4333** |
| **male dsxM*** | **7381** |

**Fig. S17. DIPA-CRISPR mutagenesis of dsxM and yellow.** Eye phenotype (arrowhead), sgRNA target locus, and Sanger sequencing chromatogram of the mutated sequence for a dsxM mosaic F0 individual (top). Eye phenotype, hatchling phenotype showing loss of pigmentation (arrowhead), sgRNA target locus, and Sanger sequencing chromatogram for a yellow mosaic F0 individual (bottom).

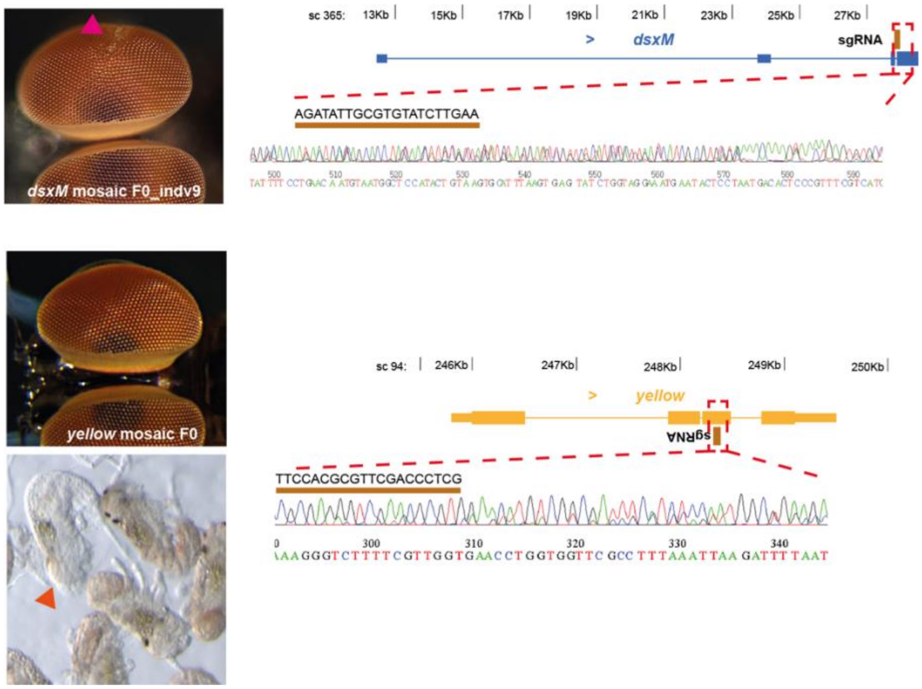

**Fig. S18. Graphical representation of the modified SCENIC pipeline used in this study.** This pipeline combines scRNA-seq with bulk ATAC-seq to identify active regulons in each cell and quantify their activity. Co-expression modules are inferred between transcription factors (TF) and genes across cells; motif discovery refines regulons by retaining targets with supporting motif evidence; cell scoring computes an AUC per cell based on the expression rank of each regulon's target genes; binarisation converts continuous AUC scores into active/inactive regulon calls across cells.

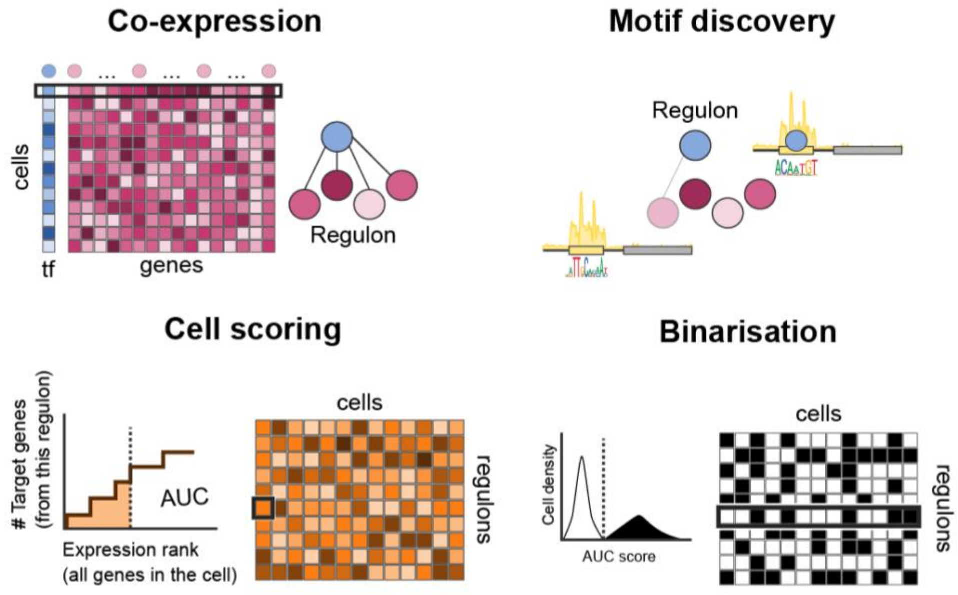

**Supplementary datasets**

**Data S1. (separate file)**

Consensus accessible putative regulatory elements (APREs) identified by ATAC-seq. One BED file per condition (NIII, NV female, NV male, NVII female, NVII male) containing IDR-filtered consensus peaks.

**Data S2. (separate file)**

SCENIC network data. Edge and node tables of the final filtered SCENIC output. For each condition, files are provided separately for the developing retina and optic lobe tissues.

**Data S3. (separate file)**

ANANSE filtered network data. Edge and node tables of the probability threshold-filtered TF–target interactions.
