## Extended methods for "A sex-specific regulator expands the embryonic ocular field to generate a novel visual system"

### Single-cell RNA-seq library preparation

Tissues were fixed in ACME solution (*1*), mechanically dissociated, and filtered to obtain a single-cell suspension, with all steps performed on ice unless otherwise stated. The four-day SPLiT-seq workflow was carried out as previously described (*1*, *2*). Briefly, on Day 1, dissociated cells were stained with Alexa Fluor 488–conjugated concanavalin A for cytoplasmic labelling and Draq5 for nuclear staining, incubated at 4 °C, and analysed by flow cytometry to quantify singlet events and assess cell-cycle phase distribution. Three successive rounds of in-cell barcoding were performed, beginning with reverse transcription using anchored oligo(dT) primers to reduce ribosomal RNA reads, followed by two ligation-based barcoding steps. On Day 2, barcoded cells were sorted by fluorescence-activated cell sorting (FACS) into lysis buffer containing Tris-HCl, NaCl, EDTA, and SDS, treated with Proteinase K, and stored at –80 °C. Day 3 comprised cDNA purification with streptavidin-conjugated magnetic beads, template switching to generate double-stranded cDNA, and PCR pre-amplification, with quantitative PCR used to determine the optimal number of cycles to remain within the exponential amplification phase. On Day 4, cDNA underwent dual size selection using Kapa Pure Beads to remove small fragments, tagmentation with the Nextera DNA Library Preparation Kit, a fourth round of PCR barcoding, and final size selection. Library concentration and fragment size distribution were assessed using a Qubit fluorometer and Agilent 2100 Bioanalyzer.

### Transcription factor annotation pipeline

To annotate transcription factors (TF), we used the proteome of the Clodip_v4 gene models (*3*). In the absence of a curated *C. dipterum* TF resource, we combined three independent evidence levels to nominate TF candidates.

1. Level 1 (protein domains) used two complementary domain-based approaches anchored to AnimalTFDB 4.0 (*4*): InterProScan (*5*) domain calls filtered against the full AnimalTFDB Pfam domain whitelist, and hmmscan against custom AnimalTFDB HMM profiles, to recover TF families poorly represented in Pfam. The AnimalTFDB Pfam domains are first described in AnimalTFDB 3.0 (*6*) and the HMM profiles in AnimalTFDB 4.0 (*4*).

2. Level 2 (GO terms) retained proteins whose GO annotation, propagated to all ancestor terms, included GO:0003700 (DNA-binding transcription factor activity) or GO:0043565 (sequence-specific DNA binding); the broader term GO:0003677 (DNA binding) was deliberately excluded as too permissive, capturing non-TF DNA-binding proteins such as histones and polymerases. This was applied separately to proteins on InterProScan (*5*) domain-based GO and full-length structure-based GO calculated with FANTASIA (*7*). We used this strategy, since some TFs lack a recognizable domain but retain informative GO transfer by homology.

3. Level 3 (homology) assessed orthology using Broccoli (*8*) and homology with protein reciprocal BLAST to a curated *D. melanogaster* TF list (AnimalTFDB (*4*)).

Independently of TF evidence levels, candidates were flagged as putative transcriptional cofactors rather than bona fide TFs if their propagated GO set contained a cofactor-associated term (e.g. GO:0003712, transcription coactivator activity; GO:0008134, TF binding) and a nuclear localization term (GO:0005634 or descendant), and lacked any direct DNA-binding TF term (GO:0003700, GO:0043565); this exclusion step was necessary because several TF-diagnostic GO terms overlap with the broader cofactor vocabulary. A parallel homology-based cofactor flag used the AnimalTFDB *D. melanogaster* cofactor list under the same orthology/homology logic as Level 3.

All proteins passing at least one evidence level (n = 1,460) were compiled into a single evidence table, recording, per protein, which sub-levels were supported, and the number of independent levels reached (Table S14). Given the pool size, candidates were triaged rather than manually curated individually. Proteins supported by only Level 1 or only Level 2 evidence were discarded outright as insufficiently corroborated. Among the remaining candidates, those reaching two or three evidence levels, or supported by Level 3 homology alone, the triage outcome depended on flagging status: unflagged proteins were accepted directly; proteins flagged only by the GO-based cofactor criterion were sent to manual curation; and proteins flagged by the homology-based cofactor criterion were discarded. Proteins sent to manual curation were checked against the literature for any evidence contradicting their classification as TFs and were rejected if such evidence was found. Accepted and manually curated candidates were combined into the final TF list used in downstream regulon analysis.

### Gene regulatory network inference using SCENIC

Gene regulatory network reconstruction was performed using the SCENIC framework (*9*, *10*), implemented in pySCENIC v0.12.1 and executed within the `aertslab/pyscenic:0.12.1` Docker image under Singularity v3.11.0 to ensure reproducibility across computational environments. The Singularity image (`aertslab-pyscenic-0.12.1.sif`) was built from Docker Hub (https://hub.docker.com/r/aertslab/pyscenic) using:

```
singularity build aertslab-pyscenic-0.12.1.sif
docker://aertslab/pyscenic:0.12.1
```

Input files preparation

*List of transcription factor*

The scenic pipeline requires a list of TF. Inspection of the transcription factor list curated by the SCENIC authors in humans, mouse and flies reveal that this list not only includes DNA binding proteins but also cofactors. This more enriched universe can reconstruct connected networks. Therefore, a SCENIC species-specific transcription factor reference for *C. dipterum* was generated using two strategies:

Identification of DNA-binding proteins. Using InterProScan (*5*) we identified the proteins that contain a DNA binding domain.

Identity to previously described transcription factors. Using the curated lists available at https://resources.aertslab.org/cistarget/ we found new candidates combining two strategies: orthology-based annotations with *Drosophila melanogaster*, *Homo sapiens*, and *Mus musculus* identified using Broccoli (*8*) orthogroups, and additional homologous matches identified by BLASTp (*11*) searches against UniProt proteomes (*12*).

*Ranking databases*

Condition-specific cisTarget ranking databases were constructed to restrict motif enrichment to regulatory regions accessible in the relevant biological context. The SCENIC analysis was performed as described with the following adjustments. Instead of constructing whole genome rankings, we only use the accessible chromatin regions. For that reason, we construct a condition-specific database.

For each condition, ATAC-seq consensus peaks from matching developmental stages were assigned to nearby genes and extracted as FASTA sequences using `cisreg_map.py` (https://github.com/m-rossello/GeneRegLocator). The resulting APREs sequences were scored against the Aerts Lab motif collection (v10nr_clust) using the `create_cisTarget_databases` pipeline (https://github.com/aertslab/create_cisTarget_databases), producing motif–gene ranking files.

*Motif annotations*

A custom motif-to-TF annotation table was generated by mapping motifs to *C. dipterum* TFs via orthology and homology, starting from *D. melanogaster* motif2TF annotations obtained from the Aerts Lab resource file `motifs-v10nr_clust-nr.flybase-m0.001-o0.0.tbl` (https://resources.aertslab.org/cistarget/motif2tf/), and replacing species-specific identifiers with the appropriate *C. dipterum* orthologs. In this table we also annotated the identity of each transcription factor from the original species to mayfly.

### SCENIC GRN inference workflow

The SCENIC workflow comprised three main steps (Fig. S17):

- GRN inference. TF-target co-expression modules were inferred from the expression matrix using the GRNBoost2 algorithm (`pyscenic grn`). This analysis was run separately for each developmental stage, with each stage-specific dataset including cells from both males and females, so that the resulting GRNs were comparable within each stage.

- Motif enrichment and regulon pruning. Modules were tested for enrichment of TF-binding motifs (`pyscenic ctx`) using the condition-specific cisTarget databases and the custom motif-to-TF annotations. The `--mask_dropouts` flag was applied to mitigate zero-inflation bias in sparse single-cell data. To maximise sensitivity, modules were not pruned at this stage (`--no_pruning`), followed by filtering based on Normalised Enrichment Score (NES), motif AUC, orthology identity, and target gene counts using create_regulons.py (https://github.com/m-rossello/SCENIC-utils).

- Regulon activity scoring. The final curated regulons were scored in each cell using the AUCell algorithm (`pyscenic aucell`) to quantify regulon activity (AUC values) and produce binarized activity matrices. Binarization thresholds were diagnosed per regulon using the custom script `diagnose_binarization.py` (https://github.com/m-rossello/SCENIC-utils), flagging regulons whose default GMM threshold might not reflect a genuine bimodal split. Each flagged regulon was inspected manually against its AUC distribution. Binarized activity matrices were then recalculated using the corrected thresholds.

Raw AUC values are not directly comparable across regulons, since different regulons have different thresholds and AUC ranges. To allow comparison, each regulon's AUC values were rescaled to a normalised activity score, using the regulon's own binarization threshold as the zero point and its maximum AUC value as the upper bound:

$$\text{normalized\_activity} = \frac{\text{AUC} - \text{threshold}}{\text{max\_AUC} - \text{threshold}}$$

Under this rescaling, a value of 0 corresponds to the activity threshold (the boundary between "on" and "off"), and a value of 1 corresponds to the most active cell for that regulon.

Custom scripts were used to patch motif annotations and apply biologically informed filtering thresholds (https://github.com/m-rossello/SCENIC-utils). The resulting regulon activity and binarized matrices were integrated with single-cell metadata in AnnData format for downstream comparative analyses. The full pipeline scripts are available at [GitHub repository link] and Zenodo, DOI: XXX.

Network topology analysis: hubs, effectors, and initiators

Directed gene regulatory networks were built from the SCENIC edge and node tables for each condition analysed. Self-loops were removed before any topology calculation. Nodes are classified as either transcription factors or target genes. Using these networks, we defined different metrics to analyse the network topology.

To understand how the nodes were connected inside the network, hub and authority scores were calculated using the HITS algorithm (*13*). Briefly, hub score measures the capacity of a node to regulate highly influential nodes. On the other hand, authority score measures how influential are the upstream nodes that regulates it. Both index are defined recursively and are interconnected, a node's hub score reflects how strongly it points to nodes with high authority, and a node's authority score reflects how strongly it is pointed to by nodes with high hub score. Hub scores were restricted to TF nodes and authority scores to target nodes, since only TFs can act as regulators and only targets can be regulated in this network.

Because both score distributions are strongly right skewed, raw scores were log transformed and standardised (z-score) across nodes with a nonzero score only. Under this normalisation, a score of zero represents average connectivity among nodes that participate in the network at all, rather than among all nodes including those with no connections. TFs with a standardised hub score of zero or above were classified as hubs. Biologically, a hub is interpreted as a regulator whose connectivity is at or above average among active TFs; it sits at the top of the regulatory hierarchy and coordinates downstream programmes rather than merely participating in them. Target genes with a standardised authority score of zero or above were classified as effectors. Biologically, an effector receives convergent, above average regulatory input from multiple upstream TFs, marking it as a terminal node where regulatory signals are integrated into transcriptional output.

A novel criterion, termed initiator, was additionally defined: a TF was classified as an initiator if zero in-degree was observed after self-loop removal, indicating that it was not itself targeted by any other regulator in the network, and is highly influential in the network. Topologically, this identifies TFs sitting at the root of the regulatory hierarchy; the entry points from which regulatory cascades originate, rather than intermediate or downstream nodes, and can reach nodes with high hub score. For each initiator, an initiator score was calculated to capture its downstream reach and influence:

$$\text{initiator\_score(i)} = \text{hub(i)} + \sum_{j} \frac{\text{hub(j)}}{\text{d(i, j)}}$$

The sum runs over all nodes j reachable from i by a directed path, d(i,j) is the length of the shortest directed path from i to j, and hub(j) is the raw, non-standardised hub score of j (zero for nodes that are not TFs or not classified as hubs). This formulation treats an initiator's influence as the sum of the hub scores of everything it can reach, discounted by path length, so that TFs feeding directly into other well-connected hubs, score higher than those whose influence is only reachable through long regulatory chains. The initiator's own hub score is added to this sum so that initiators that are simultaneously hubs are not penalised relative to initiators that only act as entry points. TFs with nonzero in degree, are assigned an initiator score of zero by definition.

Raw initiator scores were ranked normalised among initiators with a nonzero score, dividing each rank by the total number of such initiators. This rescales scores to the interval between zero and one and allows comparison across conditions with different network sizes. TFs with a normalised initiator score above 0.5, meaning those ranked in the upper half of initiators by downstream influence, were classified as initiators.

12. The UniProt Consortium, A. Bateman, M.-J. Martin, S. Orchard, M. Magrane, A. Adesina, S. Ahmad, E. H. Bowler-Barnett, H. Bye-A-Jee, D. Carpentier, P. Denny, J. Fan, P. Garmiri, L. J. D. C. Gonzales, A. Hussein, A. Ignatchenko, G. Insana, R. Ishtiaq, V. Joshi, D. Jyothi, S. Kandasaamy, A. Lock, A. Luciani, J. Luo, Y. Lussi, J. S. M.

Marin, P. Raposo, D. L. Rice, R. Santos, E. Speretta, J. Stephenson, P. Totoo, N. Tyagi, N. Urakova, P. Vasudev, K. Warner, S. Wijerathne, C. W.-H. Yu, R. Zaru, A. J. Bridge, L. Aimo, G. Argoud-Puy, A. H. Auchincloss, K. B. Axelsen, P. Bansal, D. Baratin, T. M. Batista Neto, M.-C. Blatter, J. T. Bolleman, E. Boutet, L. Breuza, B. C. Gil, C. Casals-Casas, K. C. Echioukh, E. Coudert, B. Cuche, E. De Castro, A. Estreicher, M. L. Famiglietti, M. Feuermann, E. Gasteiger, P. Gaudet, S. Gehant, V. Gerritsen, A. Gos, N. Gruaz, C. Hulo, N. Hyka-Nouspikel, F. Jungo, A. Kerhornou, P. L. Mercier, D. Lieberherr, P. Masson, A. Morgat, S. Paesano, I. Pedruzzi, S. Pilbout, L. Pourcel, S. Poux, M. Pozzato, M. Pruess, N. Redaschi, C. Rivoire, C. J. A. Sigrist, K. Sonesson, S. Sundaram, A. Sveshnikova, C. H. Wu, C. N. Arighi, C. Chen, Y. Chen, H. Huang, K. Laiho, M. Lehvaslaiho, P. McGarvey, D. A. Natale, K. Ross, C. R. Vinayaka, Y. Wang, J. Zhang, UniProt: the Universal Protein Knowledgebase in 2025. *Nucleic Acids Research* **53**, D609–D617 (2025).

13. J. M. Kleinberg, Authoritative sources in a hyperlinked environment. *J. ACM* **46**, 604–632 (1999).
